# The *Legionella* phosphocholinase AnkX modulates IMPDH2 regulation and cytoophidia dynamics

**DOI:** 10.64898/2026.01.26.701698

**Authors:** Pia Stauffer, Marietta S. Kaspers, A. Leoni Swart, Philipp Ochtrop, Bente Siebels, Vivian Pogenberg, Daniel Ferreiro Otero, Andreas Brockmeyer, Camille Schmid, Petra Janning, Hartmut Schlüter, Christian Hedberg, Aymelt Itzen, Hubert Hilbi

**Affiliations:** Institute of Medical Microbiology, University of Zürich; Zürich, Switzerland; Department of Biochemistry and Signal Transduction, University Medical Centre Hamburg Eppendorf (UKE); Hamburg, Germany; Chemical Biology Center (KBC), Department of Chemistry, Umeå University; Umeå, Sweden; Core Facility Mass Spectrometric Proteomics, University Medical Centre Hamburg Eppendorf (UKE); Hamburg, Germany; Department of Chemical Biology, Max Planck Institute of Molecular Physiology, Dortmund, Germany; Center of Structural Systems Biology (CSSB), Department of Biochemistry and Signal Transduction, University Medical Center Hamburg-Eppendorf (UKE), 20246, Hamburg, Germany

**Keywords:** cytoophidia, effector protein, GTP, host-pathogen interaction, intracellular pathogens, *Legionella pneumophila*, macrophage, metabolism, phosphocholination, post-translational modification

## Abstract

*Legionella pneumophila*, the causative agent of Legionnaires’ disease, secretes more than 300 different effector proteins into the host cell to modulate processes such as signal transduction, membrane dynamics, and metabolism. A key regulator of cellular metabolism and proliferation is inosine 5’-monophosphate dehydrogenase type II (IMPDH2), which catalyzes the rate-limiting step of *de novo* GTP biosynthesis. IMPDH2 assembles into filaments and larger structures known as cytoophidia, which enable fine-tuned regulation of enzymatic activity in response to changing nucleotide demands. Here, we identify human IMPDH2 as a previously unrecognized host target of the *Legionella* effector AnkX. We show that AnkX post-translationally modifies IMPDH2 within its regulatory domain by covalently attaching a phosphocholine moiety. While this post-translational modification does not alter the catalytic activity of IMPDH2, it disrupts filament formation, thereby impairing nucleotide-dependent regulation of enzyme activity. Consequently, AnkX-mediated phosphocholination affects IMPDH2 filament assembly, cytoophidia formation, and subcellular localization to the specific *Legionella*-containing vacuole (LCV). Thus, *L. pneumophila* subverts host GTP metabolism by posttranslationally modifying a central enzyme of the GTP biosynthetic pathway.

**Graphical abstract description:** 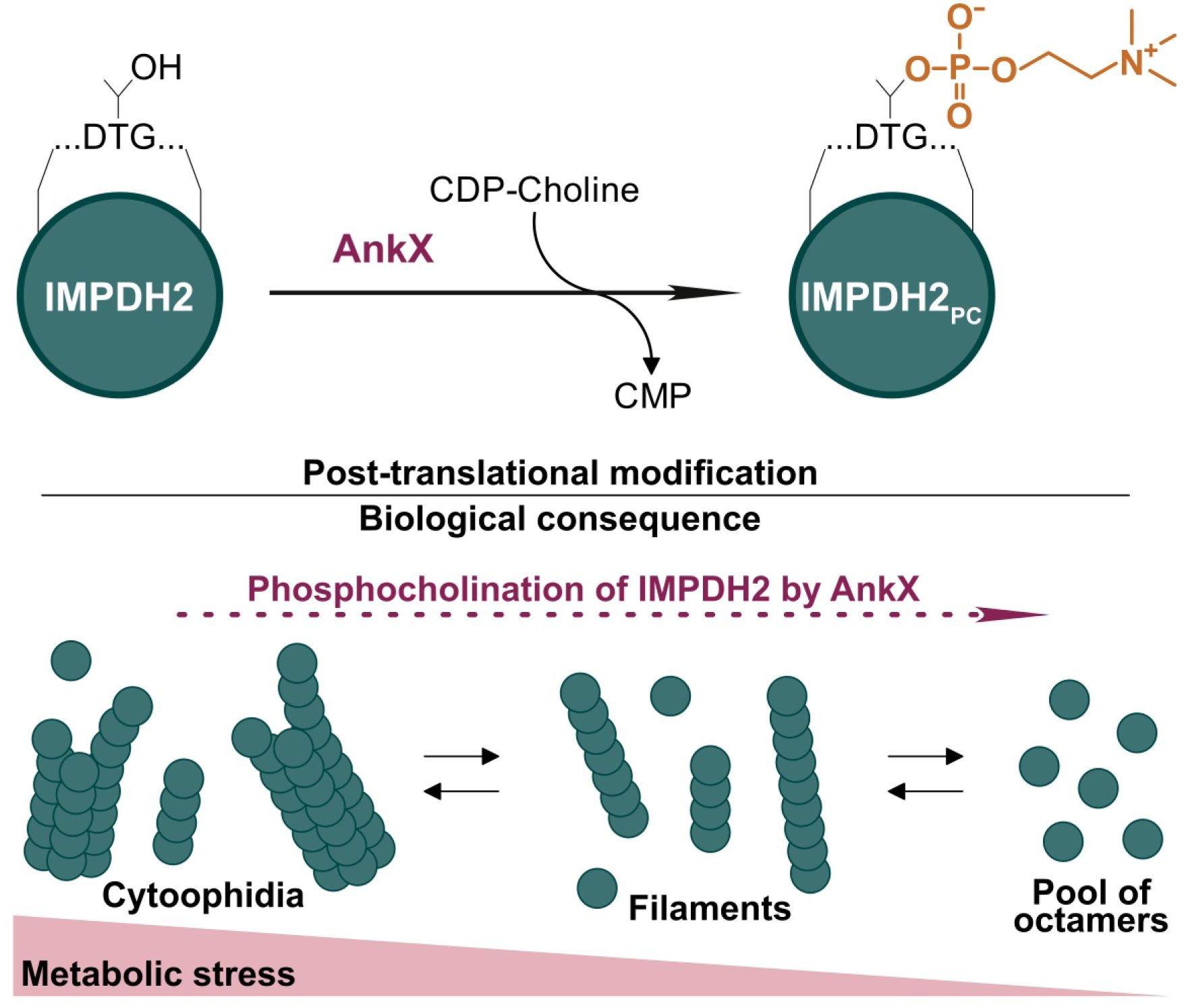

The *Legionella* effector AnkX post-translationally modifies the human host enzyme IMPDH2 by transferring a phosphocholine group using CDP-choline as co-substrate. We identified the modification site and discovered that this post-translational modification disrupts the ability of IMPDH2 to assemble into filaments and cytoophidia. By shifting the equilibrium toward free octamers, AnkX-mediated PCylation perturbs the structural regulation of IMPDH2 and alters GTP metabolism in *L. pneumophila*-infected host cells. This figure was inspired by (https://www.biorxiv.org/content/10.1101/2024.07.29.605679v2).

## Introduction

*Legionella pneumophila* is an amoebae-resistant environmental bacterium, which upon inhalation infects and destroys human alveolar macrophages, thus causing a life-threatening pneumonia called Legionnaires’ disease ^1–4^. The pivotal virulence factor of *L. pneumophila* is a type IV secretion system (T4SS) termed Icm/Dot (Intracellular multiplication/defective organelle trafficking) that translocates as many as 300 different “effector” proteins into host cells, where they subvert the cellular physiology in favor of the pathogen ^5–7^.

Some *L. pneumophila* effectors catalyze novel cell biological reactions, e.g., RidL inhibits retrograde trafficking by binding to the Vps29 retromer coat complex ^8–11^, SidC ^12,13^ and SidM/DrrA ^14–18^ bind the phosphoinositide lipid PtdIns(4)*P* with unique domains, SidM/DrrA also promotes the ATP-dependent AMPylation of the small GTPase Rab1 ^19^, and SidE family members ubiquitinate small Rab GTPases via ADP-ribosylation ^20–22^. Furthermore, the effector AnkX phosphocholinates Rab1 as well as Rab35 using the co-substrate CDP-choline, and this covalent post-translational modification is reversed by the bacterial dephosphocholinase Lem3^23–26^.

A hallmark of *L. pneumophila* virulence – conserved upon infection of macrophages or amoebae – is the formation of the replication-permissive *Legionella*-containing vacuole (LCV) ^27–29^. Formation of the ER-associated LCV is a critical outcome of *L. pneumophila*-host cell interactions, along with the subversion of a multitude of physiological processes, such as signal transduction, vesicle trafficking, membrane and cytoskeleton dynamics, as well as mitochondrial function and metabolism ^4,27,28,30^. Proteomics analyses of intact LCVs isolated from macrophages or *Dictyostelium discoideum* amoebae ^31–34^ implicated the small GTPase Ran ^35,36^ and the large fusion GTPase atlastin/Sey1 ^37–39^ in LCV formation. Moreover, inosine 5’-monophosphate (IMP) dehydrogenase (IMPDH) was also identified by proteomics on LCVs isolated from *L. pneumophila*-infected macrophages ^32^.

IMPDH catalyses the first and rate-limiting step of the *de novo* GTP synthesis pathway, the NAD⁺-dependent conversion of IMP to xanthine 5’-monophosphate (XMP) and thus is a central regulator of the intracellular GTP levels and cell proliferation ^40–42^. IMPDH exists in mammalian cells as two isoforms, IMPDH1 and IMPDH2, which share 84% sequence identity^43^. In most tissues, IMPDH1 is constitutively expressed at basal levels, while IMPDH2 is the dominant isoform. An upregulation of IMPDH2 expression is associated with proliferating and cancerous cells ^44–46^.

The GTP and ATP *de novo* biosynthesis pathways are linked via their common precursor IMP, and adenine and guanine nucleotides as well as IMP allosterically regulate IMPDH and the balance between ATP and GTP pools in the cell ^47–49^ (**Fig. S1a**). The catalytic activity and subcellular distribution of IMPDH are tightly regulated on multiple levels. IMPDH monomers consist of a catalytic and a regulatory domain (called Bateman domain) and assemble into tetramers that reversibly dimerize to form octamers. While binding of ATP or ADP to the Bateman domain of each monomer induces an extended octamer conformation and increased catalytic activity, binding of GTP or GDP leads to a compressed conformation and enzyme inhibition ^48^ (**Fig. S1b**). In addition to allosteric regulation, IMPDH activity is regulated by post-translational modifications, splice variants, and polymerization into filaments, all of which alter the enzyme’s resistance against allosteric inhibition by guanine nucleotides ^48,50–53^.

Polymerization of IMPDH2 octamers into filaments is induced by ATP or GTP binding ^54,55^. In cells, the assembly of IMPDH2 filaments into large fiber-like bundles, termed cytoophidia, occurs upon increased GTP demand, guanine nucleotide deprivation, or pharmacological inhibition, and is facilitated by molecular crowding ^48,50,56–60^ (**Fig. S1c**). Assembly into cytoophidia prolongs the half-life of IMPDH2 and allows purine nucleotide pools to expand under conditions of high demand ^50,60,61^. Thus, the ability of IMPDH2 to assemble into cytoophidia is instrumental for the fine-tuning of purine nucleotide homeostasis under physiological and pathological conditions such as embryonic development, cancer, and neurodegeneration ^62–64^. However, the molecular mechanisms orchestrating cytoophidia dynamics remain elusive.

In this study, we identify IMPDH2 as a phosphocholination substrate of the *L. pneumophila* effector AnkX (**Fig. S1d**). Phosphocholination of Thr147 is a novel post-translational modification of IMPDH2, which modulates the nucleotide-dependent regulation of enzyme activity, oligomerization state, and subcellular localization at LCVs. Accordingly, *L. pneumophila* augments intracellular replication by targeting a pivotal enzyme in the *de novo* GTP biosynthesis pathway.

## Results

### *Legionella* effector AnkX phosphocholinates IMPDH2 and modulates its regulation

In addition to its natural co-substrate, CDP-choline, the *L. pneumophila* effector AnkX can use synthetic CDP-choline derivatives as co-substrates for the post-translational modification of target proteins ^65^. To detect and identify additional targets of AnkX, we employed a biotin-functionalized CDP-choline derivative (CDP-choline-biotin, CDP-CB) to enzymatically modify putative substrate proteins with phosphorylcholine-biotin (**Fig. 1a**). Western blot analysis of HeLa cell lysate co-incubated with AnkX and CDP-CB revealed three modified protein species at approximately 26, 60 and 95 kDa, respectively, which were absent in the control sample without AnkX (**Fig. 1b**). To identify these phosphorylcholine-biotinylated proteins, they were enriched with streptavidin-coupled magnetic beads, separated by SDS-PAGE, and analyzed by LC-MS/MS. Thus, the individual bands at 26 kDa were identified as Rab proteins (Rab1a, Rab1b, Rab2a, Rab2b, Rab30, Rab35, and Rab43), at 95 kDa as AnkX, and at 60 kDa as IMPDH2. Western blot analysis using an IMPDH2-specific antibody further confirmed the presence of IMPDH2 (**Fig. 1c**).

**Figure 1.**
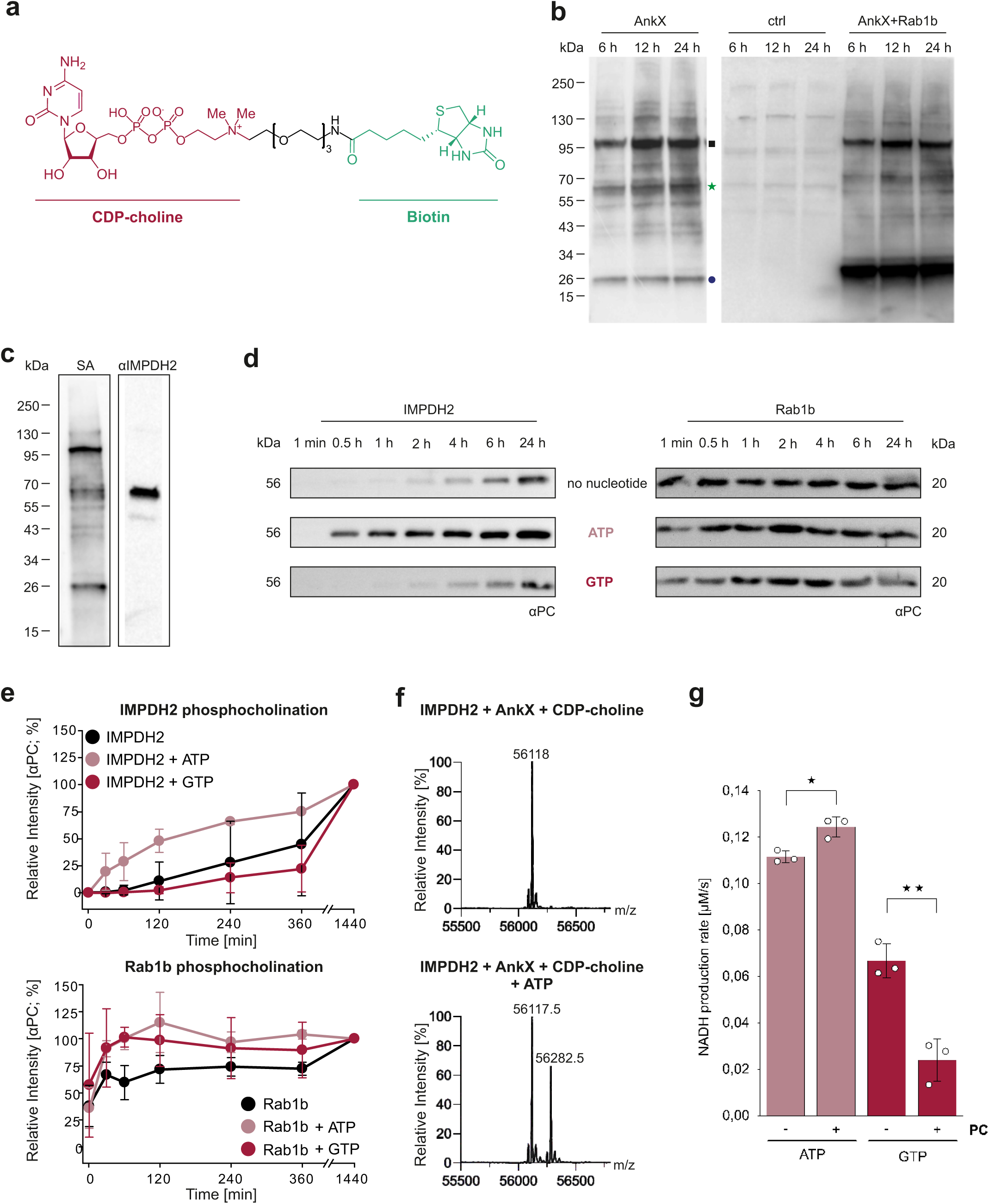
AnkX phosphocholinates IMPDH2 preferably in presence of ATP and enhances negative feedback inhibition of IMPDH2. (**a**) Structural formula of CDP-choline-biotin (CDP-choline: red, linker: black, biotin: green). (**b**) HeLa cell lysate was incubated for 6, 12, or 24 h with 100 µM CDP-choline-biotin (ctrl) or CDP-choline-biotin and 1 µM AnkX. As a positive control, samples were additionally supplemented with Rab1b. Labelled target proteins were separated by SDS-PAGE and detected by Western blot using a streptavidin-fluorescein conjugate. AnkX (black square), IMPDH2 (green star), Rab GTPases (blue circle). (**c**) Biotinylated target proteins in HeLa lysate were enriched by pulldown with streptavidin-coupled magnetic beads and separated by SDS-PAGE for analysis of individual bands by tandem mass spectrometry (MS/MS). The enrichment of IMPDH2 was verified by Western blot analysis with streptavidin (SA) and a specific anti-IMPDH2 antibody (αIMPDH2). (**d**) Phosphocholination kinetics of purified IMPDH2 or Rab1b (5 µM) upon addition of AnkX (0.5 µM) and CDP-choline (100 µM) in absence or presence of 100 µM ATP or GTP were assessed by Western blot with an anti-phosphocholine antibody (αPC), followed by a horseradish peroxidase (HRP)-coupled antibody and chemiluminescence. (**e**) Quantification of (d) by densitometry. (**f**) IMPDH2 (5 µM) was incubated with ATP (1 mM, 1 h) or not, followed by CDP-choline (100 µM, 2h) and AnkX (0.5 µM, 2 h), and phosphocholination was assessed by intact mass spectrometry iMS). (**g**) IMPDH2 activity (NAD⁺-dependent conversion of IMP to XMP) was quantified measuring NADH production over time for phosphocholinated and unmodified IMPDH2 in presence of ATP or GTP. Means (±SD) represent three independent biological replicates (unpaired, two-tailed t-test; *, p < 0.05; **, p < 0.01).

To validate the ability of AnkX to phosphocholinate IMPDH2 *in vitro* and assess the kinetics of IMPDH2 phosphocholination, we incubated recombinantly produced and purified IMPDH2 with AnkX and CDP-choline. The phosphocholination reaction was performed in presence or absence of ATP or GTP and quantified by anti-phosphocholine Western blot (**Figs. 1d, S2**). Under all conditions tested, we observed a time-dependent phosphocholination of IMPDH2. However, IMPDH2 was only minimally modified in the absence of nucleotides or in presence of GTP, whereas the presence of ATP significantly increased IMPDH2 phosphocholination (**Fig. 1e**). This ATP-dependent phosphocholination was specific for IMPDH2 and did not occur with the previously reported AnkX target Rab1b. Under the conditions tested, Rab1b was modified faster than IMPDH2 and reached full modification after 1 h. We further analyzed the phosphocholination of purified recombinant IMPDH2 by AnkX *in vitro* using intact mass spectrometry (iMS), demonstrating that IMPDH2 is modified with a single phosphocholine moiety (**Fig. 1f**).

Next, we studied the effect of phosphocholination on the enzymatic activity of IMPDH2 (NAD⁺-dependent oxidation of IMP to XMP) by monitoring NADH production over time. The enzymatic activity of phosphocholinated IMPDH2 (IMPDH2_PC_) was identical to control experiments with AnkX or CDP-choline alone (**Fig. S3a**), indicating that phosphocholination does not alter the catalytic activity of IMPDH2. The results demonstrated that IMPDH2_PC_ retains catalytic activity in the presence of ATP with turnover rates comparable to or exceeding those of the unmodified enzyme (**Figs. 1g, S3b**). As previously reported ^48^, GTP binding inhibits IMPDH2 activity, which under the conditions used was approximately 40% decreased compared to the presence of ATP. Intriguingly, however, IMPDH2_PC_ exhibited a significantly stronger activity reduction by approximately 75% in presence of GTP compared to the unmodified enzyme. Therefore, phosphocholination renders IMPDH2 more sensitive to GTP inhibition (**Figs. 1g, S3b**).

During *L. pneumophila* infection, Rab1 phosphocholination is reversed by the effector protein Lem3, which thus acts as an antagonist of AnkX ^23–26^. However, Lem3 does not dephosphocholinate IMPDH2 *in vitro* (**Fig. S3c**), indicating that Lem3 antagonizes the phosphocholination of Rab1 but not of IMPDH2.

In summary, IMPDH2 is phosphocholinated by the *L. pneumophila* effector AnkX in an ATP-dependent manner but not dephosphocholinated by Lem3. The covalent modification does not alter the enzymatic activity of IMPDH2 *per se* but sensitizes the enzyme towards inhibition by GTP.

### Phosphocholination causes disassembly of ATP-induced IMPDH2 filaments

IMPDH2 octamers reversibly oligomerize into linear filaments *in vitro* upon binding of ATP or GTP ^54,55^. To test whether phosphocholination affects the filamentation state of IMPDH2, we induced filament formation of IMPDH2 by addition of ATP, followed by incubation with CDP-choline, AnkX, or the catalytically inactive mutant AnkX_H229A_. Negative stain transmission electron microscopy (TEM) revealed that ATP-induced IMPDH2 filaments disassembled in the presence AnkX and CDP-choline (**Fig. 2a**). In control experiments, IMPDH2 filaments were not disassembled in the presence of catalytically deficient AnkX_H229A_, or AnkX or CDP-choline alone. This effect is due to IMPDH2 phosphocholination as demonstrated by parallel iMS sample analysis (**Figs. 2b**). Thus, the loss of filament formation capacity of IMPDH2 depends on AnkX-mediated phosphocholination.

**Figure 2.**
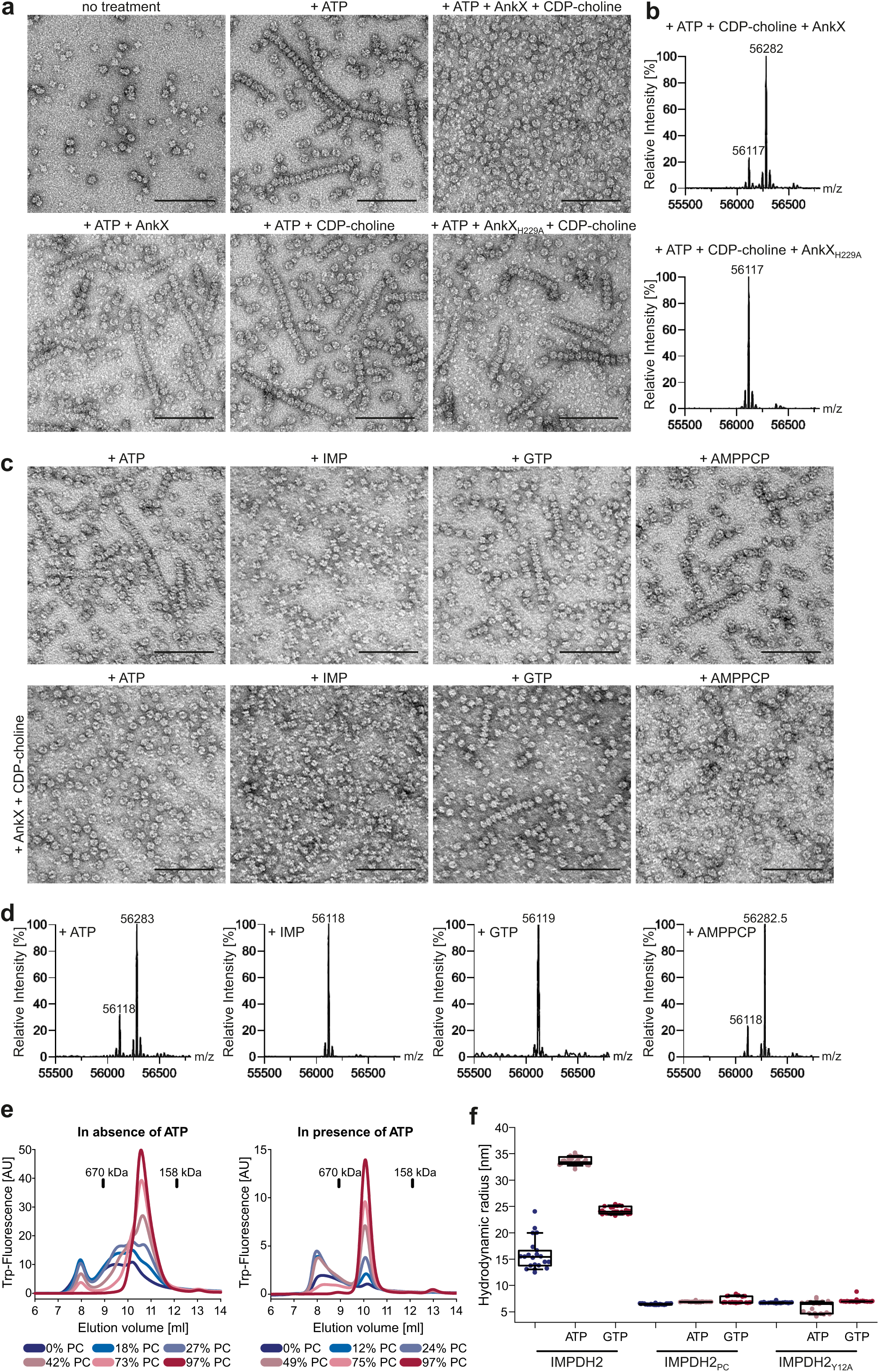
Phosphocholination of IMPDH2 causes disassembly and prevents formation of ATP-induced filaments. (**a**, **b**) Purified IMPDH2 (5 µM) was incubated with 1 mM ATP (1 h, RT) to induce filament formation or left untreated, followed by purified AnkX_1-800_ or the catalytically inactive mutant AnkX_1-800_H229A_ (0.5 µM), and/or CDP-choline (100 µM) for an additional 2 h. Samples were (**a**) analyzed by negative staining and TEM (scale bars, 100 nm), or (**b**) by MS. Phosphocholinated IMPDH2, represented as a mass difference of 165 Da, was only detected in the sample containing catalytically active AnkX_1-800_. (**c**, **d**) Purified IMPDH2 (5 µM) was treated for 1 h with either 1 mM ATP, 1 mM IMP, 1 mM GTP, or 1 mM AMPPCP and 1 mM MgCl_2_, then further incubated for 2 h with AnkX_1-800_ (0.5 µM), and CDP-choline (100 µM) or left without additional treatment. Samples were analyzed by (**c**) negative staining and TEM (scale bars, 100 nm), or (**d**) MS. (**e**, **f**) Analytical size exclusion chromatography (aSEC) of unmodified and phosphocholinated IMPDH2 in the absence or presence of ATP. Purified IMPDH2 (50 μg) phosphocholinated to varying percentages (indicated below the respective graph) was loaded and detected at 340 nm using tryptophan-fluorescence (excitation at 280 nm). Elution volumes of reference proteins (thyroglobulin: 670 kDa, γ-globulin: 158 kDa) are indicated by a vertical line. (**g**) Dynamic light scattering (DLS) was conducted on IMPDH2, IMPDH2_PC_, and IMPDH2_Y12A_, in the presence and absence of ATP and GTP as indicated. Each data point represents an individual measurement (n > 21). The boxes indicate the median (horizontal line within the box) and extend from the 25^th^ to the 75^th^ percentile, with standard deviation reaching the 90^th^ and 10^th^ percentiles.

Moreover, to directly visualize IMPDH2 filament disassembly, we employed negative stain TEM. Indeed, IMPDH2 filaments induced by ATP or the non-hydrolysable ATP analogue AMPPCP disassembled upon phosphocholination (**Fig. 2cd**). However, we observed no disassembly or phosphocholination of GTP-treated IMPDH2, and IMP did not cause filamentation. Together with the data shown in Fig. 1d this suggests that AnkX specifically phosphocholinates ATP-bound filamentous IMPDH2. Furthermore, filament breakdown is not a result of ATP-hydrolysis, since AMPPCP-bound IMPDH2-filaments also disassembled after phosphocholination.

We further assessed filament formation using analytical size exclusion chromatography (aSEC) (**Figs. 2e, S4**). To this end, quantitatively phosphocholinated IMPDH2 was analysed, along with samples modified to approximately 10, 25, 50 and 75%. These partially modified samples were generated by mixing phosphocholinated and unmodified IMPDH2 in defined ratios. The exact degree of modification was determined by iMS (**Fig. S4a**). The gradual shift in retention volume towards lower molecular weight with increasing proportions of IMPDH2_PC_ illustrates a shift in the equilibrium from filaments towards smaller species (**Figs. 2e, S4b**), in the range between 670 and 158 kDa. We observed this shift both in presence and in absence of ATP. The retention volume of all observed species shifts towards a higher molecular weight in the presence of ATP, likely representing the extended, ATP-bound conformation of IMPDH2.

To further characterize the size of IMPDH2_PC_ particles, we performed dynamic light scattering (DLS) experiments in absence or presence of ATP or GTP (**Fig. 2f**). Unmodified IMPDH2 consistently exhibited larger hydrodynamic radii compared to IMPDH2_PC_, regardless of the nucleotide-loading status. As a reference, we included the IMPDH2_Y12A_ mutant, which is defective for filament formation and has been described to predominantly adopt tetrameric and octameric states ^55^. IMPDH2_PC_ and IMPDH2_Y12A_ exhibited similar hydrodynamic radii, further illustrating the loss of filament-forming ability of IMPDH2_PC_ and suggesting the formation of tetrameric or octameric species. In summary, these data indicate that phosphocholination of IMPDH2 leads to filament disassembly, stabilizes lower order oligomers, and confers resistance towards ATP-or GTP-induced filament assembly.

### IMPDH2 is phosphocholinated in the regulatory Bateman domain

Having established the phosphocholination of IMPDH2 by AnkX, we sought to identify the specific modified amino acid of IMPDH2. Quantitatively modified IMPDH2_PC_ (**Figs. 3a, S4a**) was subjected to tryptic digestion and analysed by LC-MS/MS. We employed a combination of different fragmentation techniques, namely collision-induced dissociation (CID), higher-energy collisional dissociation (HCD) and electron transfer dissociation (ETD). During CID and HCD, the phosphocholine moiety is cleaved from the modified amino acid producing a characteristic diagnostic ion at m/z 184 which induced signal suppression. Subsequent fragmentation of this reporter ion showed the presence of characteristic fragment ions (m/z 60 ([CH3)3NH]+), m/z 86 ([MH-H3PO4]+), m/z 104 ([MH-HPO3]+), und m/z 125 ([H2PO4)(CH2)2]+)) for a peptide with m/z 532.59 (**Figs. 3b, S5ab**) ^23^. ETD analysis, suited for labile PTMs of peptides identified by LC-MS/MS ^66^, consistently identified threonine 147 (T147_IMPDH2_), the last threonine within the peptide HGFCGIPITD**T**GR (m/z 532.59, +3 charge state), as the phosphocholinated residue (**Figs. 3c, S5cd**). Taken together, tandem MS revealed that T147_IMPDH2_ located in the regulatory Bateman domain (**Fig. S1a**) is phosphocholinated by AnkX.

**Figure 3.**
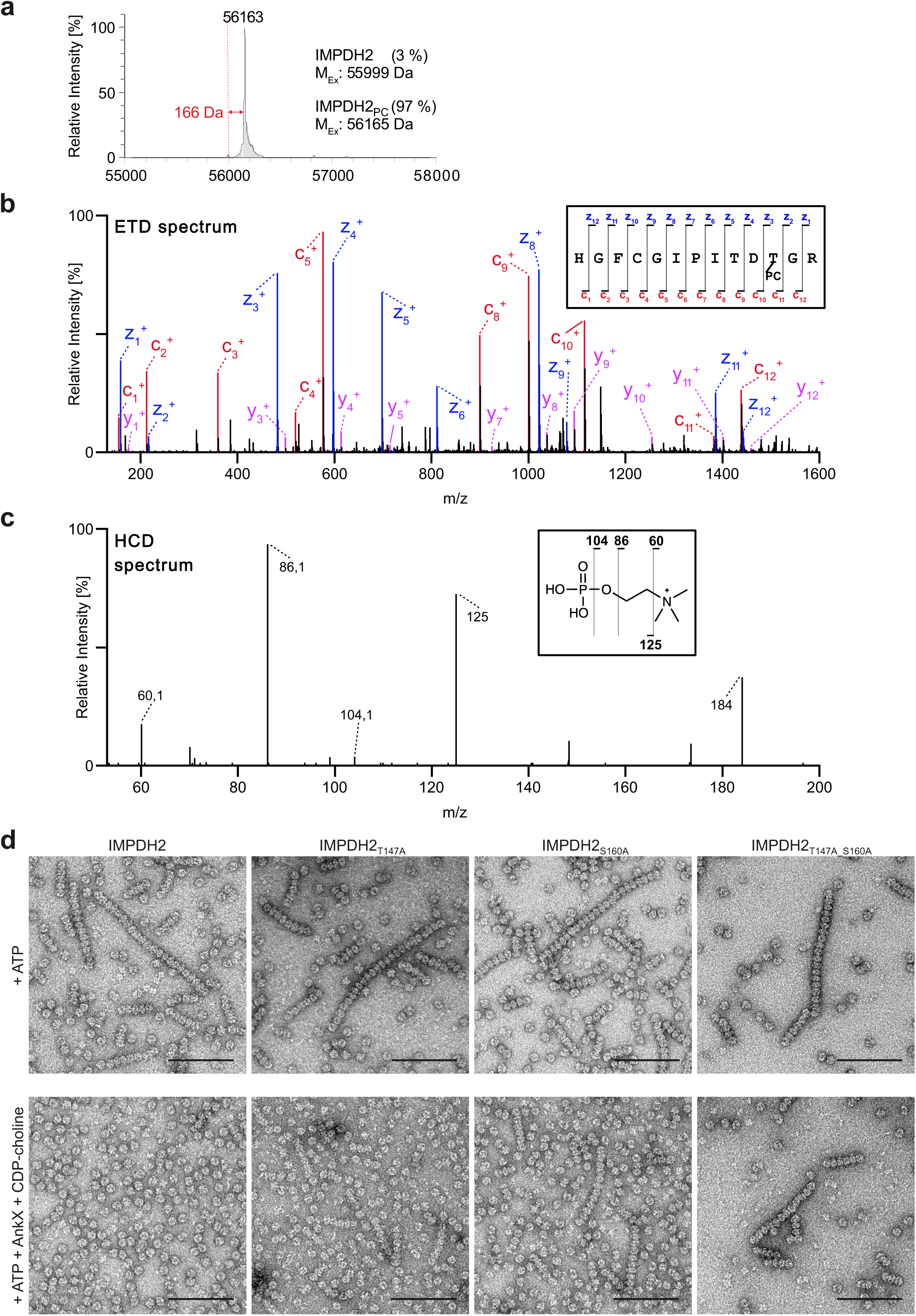
IMPDH2 is phosphocholinated in the regulatory Bateman domain. (**a-c**) Purified IMPDH2 was quantitatively phosphocholinated *in vitro* and initially analyzed by iMS to determine the percentage of phosphocholination, followed by tryptic digestion of the sample and analysis using LC-MS/MS. (**a**) iMS revealed a peak shift of 166 Da for IMPDH2_PC_ compared to unmodified IMPDH2, corresponding to a modification by a single phosphocholine moiety at a modification efficiency of 97%. (**b**) MS2 ETD fragmentation spectrum of the peptide HGFCGIPITDTGR (aa 137–149, precursor m/z 532.5854) identifying T147_IMPDH2_ as the sole phosphocholinated residue. (**c**) MS3 HCD fragmentation spectrum proves the presence of a phosphocholine. (**d**) Filament assembly of wild-type IMPDH2, IMPDH2_T147A_, IMPDH2_S160A_, and IMPDH2_T147A_S160A_ was induced by addition of 1 mM ATP for 1 h. After additional 2 h incubation with AnkX_1-800_ (0.15 µM) and CDP-choline (100 µM), filaments were assessed by negative staining TEM. Scale bars, 100 nm.

Having identified the modification site T147_IMPDH2_, we tested phosphocholination and filament (dis)assembly of the mutant IMPDH2_T147A_ (**Fig. 3d**). To this end, we treated IMPDH2_T147A_ with ATP and subsequently with AnkX and CDP-choline. IMPDH2_T147A_ formed filaments upon ATP-treatment comparable to wild-type IMPDH2 filaments. After subsequent incubation with AnkX and CDP-choline, the filaments persisted, but they were shorter than the filaments in absence of AnkX (**Fig. 3d**). Unexpectedly, IMPDH2_T147A_ was still phosphocholinated, as detected by iMS (**Fig. S5e**).

IMPDH1 harbors amino acid S160 as a phosphorylation site in the Bateman domain, and its modification affects the sensitivity of the enzyme to allosteric inhibition by GTP/GDP ^51^. The deletion of S160 in IMPDH2 renders the protein incapable of filament formation ^67^. Given the exposed position of S160, the similar properties of a phosphoryl group and a phosphocholine group, and the structural and biological implications described for S160, we also tested IMPDH2_S160A_ and the double mutant IMPDH2_T147A_S160A_ for phosphocholination (**Fig. 3d**). Both mutants showed filament formation in presence of ATP comparable to the wild-type protein. After treatment of the IMPDH2_S160A_ and IMPDH2_T147A_S160A_ filaments with AnkX and

CDP-choline, we observed filaments that appeared slightly shorter (**Fig. 3d**) but were still phosphocholinated, as detected by iMS (**Fig. S5f**). Taken together, these data indicate that T147 is the preferred phosphocholination site of IMPDH2. However, if T147 is not available, another exposed serine, threonine, or tyrosine, likely including S160, might also be phosphocholinated. All mutant proteins tested showed a defect in phosphocholination-induced filament disassembly, supporting a biological relevance of these amino acid residues.

### AnkX induces disassembly of IMPDH2 cytoophidia in *L. pneumophila*-infected cells

Oligomerization and assembly of IMPDH2 represents an important regulatory mechanism to sustain enzyme activity during high cellular GTP demand ^61^. Cells respond to starvation or treatment with the IMPDH2 inhibitors mycophenolic acid (MPA) or ribavirin with the assembly of IMPDH2 into large filamentous structures called cytoophidia ^54,56,57,68^. Since cytoophidia can be visualized by fluorescence microscopy, we explored whether AnkX-mediated IMPDH2-cytoophidia disassembly occurs during an *L. pneumophila* infection. To this end, we generated the *L. pneumophila* Δ*ankX* deletion mutant strain in the JR32 strain background and compared its effect on cellular cytoophidia with the parental strain and the type IV secretion defective mutant Δ*icmT*.

MPA-treated A549 lung epithelial cells were infected for 2 h with these GFP-producing *L. pneumophila* strains and analyzed for cytoophidia by immunofluorescence confocal microscopy staining against IMPDH2 (**Fig. 4ab**). While more than 90% of uninfected MPA-treated cells showed cytoophidia, only about 10% of wild-type *L. pneumophila*-infected cells were cytoophidia-positive (**Fig. 4b**). In contrast, Δ*ankX*-and Δ*icmT*-infected cells were >70% and >90% cytoophidia-positive, respectively, and cytoophidia disassembly was restored in the Δ*ankX* deletion mutant upon providing *ankX* on a plasmid. These findings demonstrate that *L. pneumophila* disassembles IMPDH2 cytoophidia in infected host cells, and the disassembly requires the secreted effector AnkX as well as the Icm/Dot T4SS.

**Figure 4.**
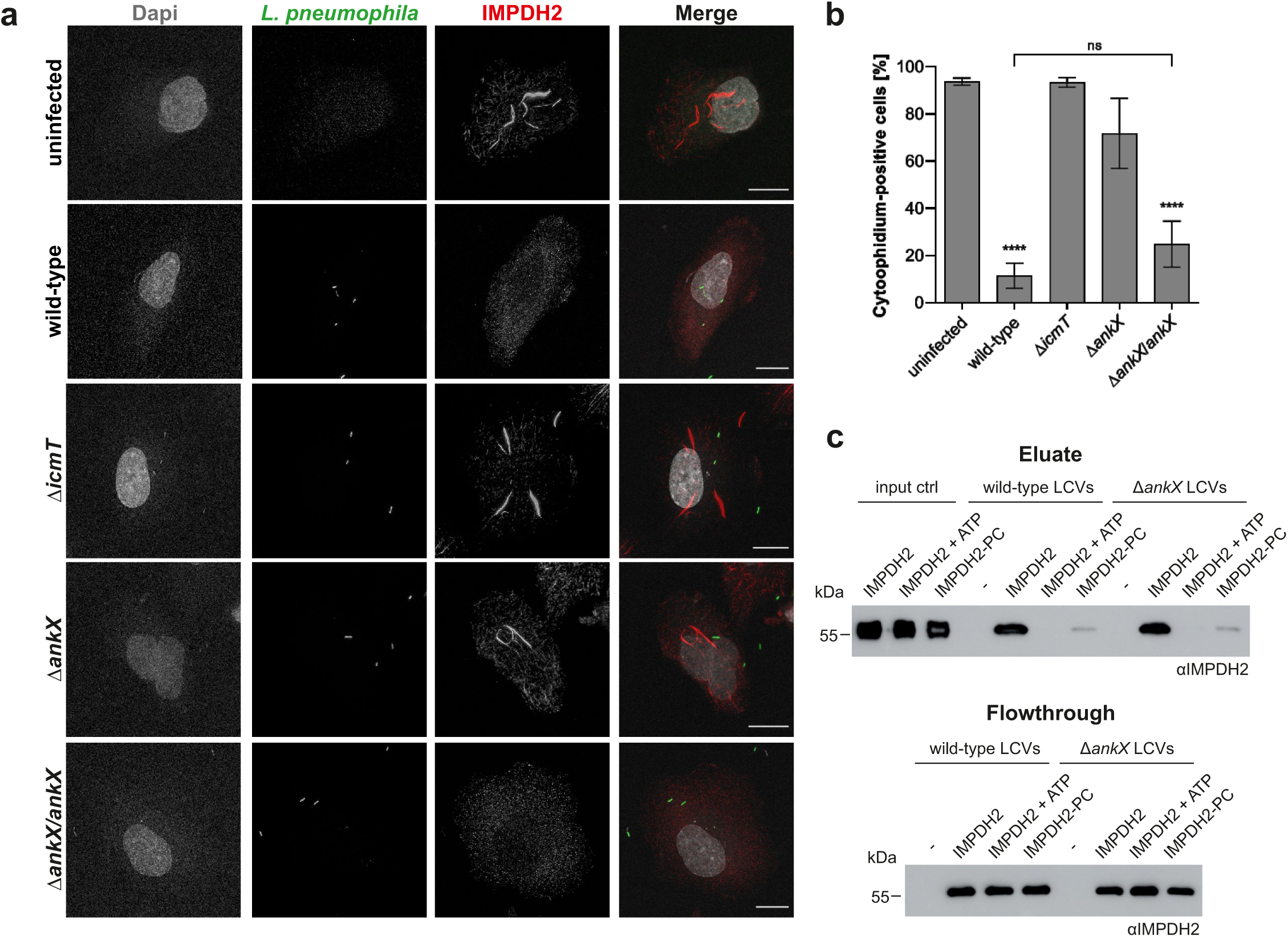
*L. pneumophila* disassembles IMPDH2 cytoophidia in the host cell. (**a**) A549 cells were treated with 500 nM IMPDH2 inhibitor MPA and infected (MOI 100, 1 h) with GFP-producing *L. pneumophila* wild-type, Δ*ankX* or Δ*icmT* (pNT28), with Δ*ankX*/*ankX* (pLS284), or left uninfected. Cells were washed, and fresh medium with 500 nM MPA was added for another 2 h, fixed with 4% PFA, stained for IMPDH2 (red) and DAPI (grey), and analyzed by confocal microscopy. Images are maximum intensity projections. Scale bars, 10 µm. (**b**) The percentage of cytoophidia-positive cells was scored (n > 800/strain, 3 independent experiments, one-way ANOVA with Tukey’s multiple comparisons test; ****, P<0.0001). (**c**) Murine RAW 264.7 macrophages were infected (MOI 50, 1 h) with *L. pneumophila* wild-type or Δ*ankX*, washed and homogenized, and LCVs were isolated by immuno-magnetic separation. Purified IMPDH2 (5 µM) pre-incubated for 6 h at 25°C with 1 mM ATP (IMPDH2 + ATP), or ATP, 0.5 µM AnkX_1-800_, and 100 µM CDP-choline (IMPDH2-PC), or none (IMPDH2), was diluted to 1 µM and added to the LCV samples (15 min, RT). As a negative control, LCVs were incubated with buffer only (-). LCVs were washed, eluted, and protein binding was analyzed by anti-IMPDH2 Western blot (upper panel). Flow-through samples were collected after incubation of LCVs to verify equal passage of filamentous ATP-bound and unmodified IMPDH2 through the immuno-magnetic separation columns (lower panels).

### AnkX and IMPDH2 localize to LCVs

AnkX localizes to the plasma membrane, as well as to vesicular and tubular structures and LCVs through binding to phosphatidylinositol-3-phosphate (PtdIns(3)*P*) and PtdIns(4)*P*, where it interacts with the targets Rab1 and Rab35 ^69,70^. We established the binding of purified His_6_-AnkX and its catalytically inactive mutant His_6_-AnkX_H229_ to isolated LCVs (**Fig. S6a**). Binding of AnkX also occurred to LCVs harboring Δ*sidM L. pneumophila*, and therefore, does not require Rab1, which is recruited by the effector SidM to LCVs. Therefore, wild-type and catalytically inactive AnkX bind to LCVs independently of the presence of Rab1.

Having confirmed the localization of AnkX to LCVs, we hypothesized that IMPDH2 binds to LCVs as well. To test this, we purified LCVs from RAW 264.7 macrophages infected with wild-type or Δ*ankX L. pneumophila* by magnetic activated cell sorting (MACS) ^32,34^. LCVs immobilized on the MACS columns were incubated with untreated IMPDH2 or IMPDH2 pre-treated with ATP in absence or presence of AnkX and CDP-choline. Subsequently, LCVs were eluted, and binding of IMPDH2 was assessed by anti-IMPDH2 Western blot (**Fig. 4c**). To rule out the possibility that the filamentation of IMPDH2 affects the passage through the column, we also analyzed the flow through by anti-IMPDH2 Western blotting (**Fig. 4c**). Using this approach, untreated IMPDH2 bound to LCVs from wild-type or Δ*ankX*-infected macrophages, confirming previous proteomics identifying IMPDH2 on LCVs isolated from macrophages ^32^. Strikingly, we observed barely any binding of ATP-treated IMPDH2 or IMPDH2_PC_, while the amount of IMPDH2 in the column flow through did not change. These results indicate that assembly into filaments prevents IMPDH2 binding to LCVs. Phosphocholination of IMPDH2 by incubation with ATP, AnkX, and CDP-choline was confirmed by anti-phosphocholine Western blot (**Fig S6b**) and resulted in the binding of low amounts of IMPDH2 to LCVs (**Figs. 4c, S6c**). In summary, AnkX binds to LCVs regardless of its catalytic activity, and untreated purified IMPDH2 but not ATP-treated IMPDH2 filaments also localizes to LCVs.

### IMPDH2 promotes intracellular replication of *L. pneumophila*

Given that *L. pneumophila* affects the post-translational modification and subcellular targeting of IMPDH2, we sought to investigate whether IMPDH2 is implicated in intracellular replication of the bacteria. To this end, we tested the effects of pharmacological inhibition of IMPDH2 on intracellular growth of GFP-producing *L. pneumophila* in infected A549 cells or RAW 264.7 macrophages (**Figs. 5a, S7a**). Treatment of host cells with increasing concentrations of MPA impaired intracellular growth of *L. pneumophila* in a dose-dependent manner. To rule out any toxic effects of MPA on *L. pneumophila*, we treated bacteria growing in liquid culture with the same MPA concentrations (**Fig. S7b**). Extracellular *L. pneumophila* growth remained unaffected by MPA except for the highest tested concentration (25 μM). We also tested whether the lack of AnkX or its antagonist Lem3 affects the intracellular growth of *L. pneumophila* in presence of MPA. Both deletion mutants, Δ*ankX* and Δ*lem3*, exhibited a slight intracellular growth defect compared to wild-type *L. pneumophila*; however, their replication was equally impaired by MPA (**Fig. 5a**).

**Figure 5.**
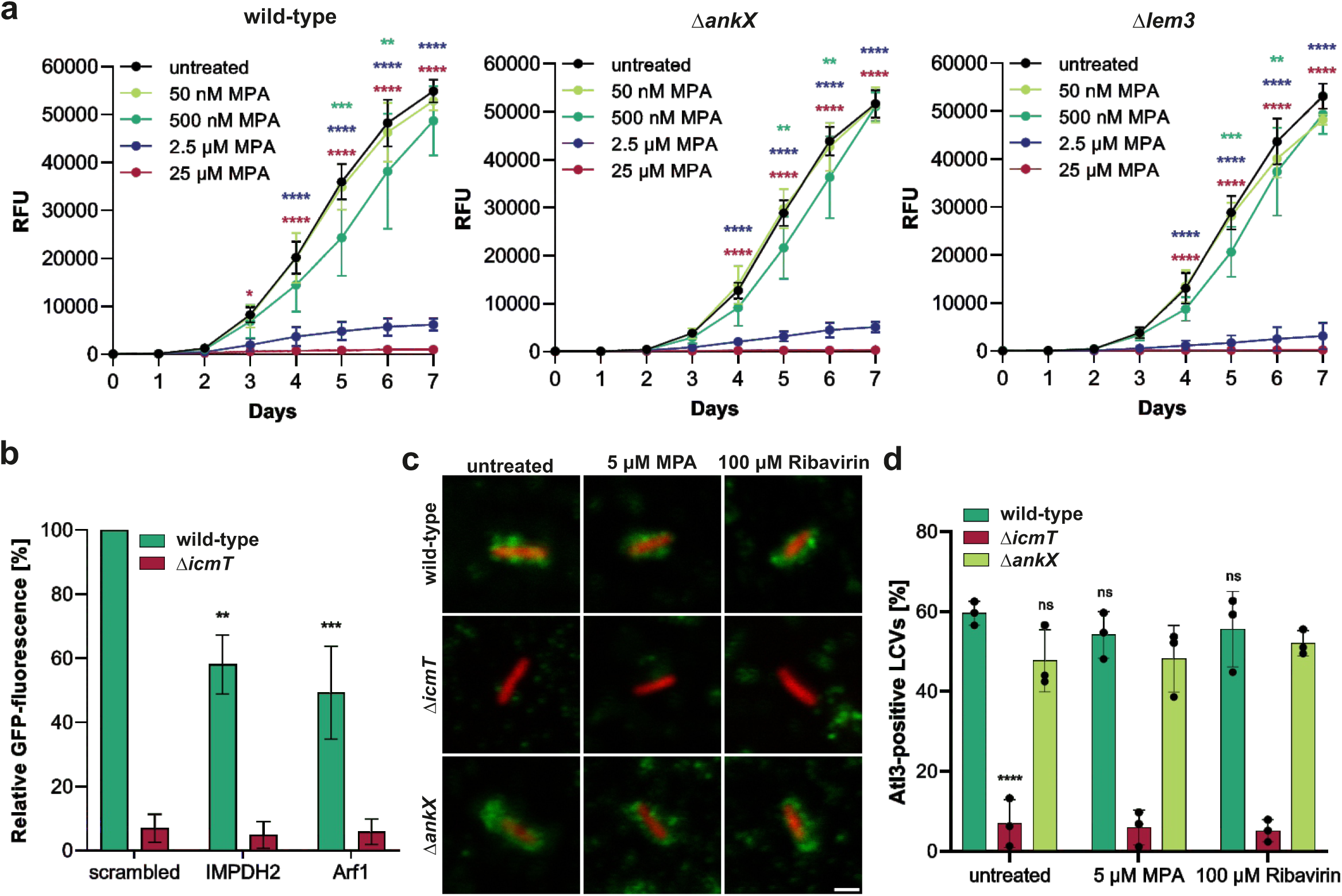
IMPDH2 promotes intracellular growth of *L. pneumophila*. (**a**) A549 cells were treated with MPA at the indicated concentrations and infected (MOI 1) with GFP-producing *L. pneumophila* wild-type, Δ*ankX,* or Δ*lem3* (pNT28), and intracellular replication at 37°C was assessed daily by relative fluorescent units (RFU). For each time point, the difference of the MPA-treated cells to the untreated cells was tested by a two-way ANOVA with Dunnett’s multiple comparisons test; **, P < 0.01; ***, P < 0.001; ****, P < 0.0001. Means and SD of three independent experiments are shown. (**b**) A549 cells were transfected for 48 h with 10 nM siRNA oligonucleotides targeting IMPDH2, or Arf1 (positive control), or with AllStars unspecific oligonucleotides (“scrambled”), infected (MOI 10) with GFP-producing *L. pneumophila* wild-type or Δ*icmT* (pNT28), and intracellular growth was assessed by fluorescence increase between 1 h and 24 h p.i. using a fluorescence plate reader. Means and SEM of results from four independent experiments are shown (two-way ANOVA with Dunnett’s multiple comparisons test; **, P < 0.01; ***, P < 0.001; all groups compared to “scrambled”, wild-type infected). (**c**) RAW 264.7 macrophages were infected (MOI 50) with mCherry-producing *L. pneumophila* wild-type, Δ*icmT,* or Δ*ankX* (pNP102) and simultaneously treated with 5 µM MPA or 100 µM ribavirin or left untreated. 2 h post-infection, cells were homogenized, LCVs were fixed with 4% PFA, stained for Atl3 (green) and analyzed by confocal microscopy. Scale bars, 1 µm. (**d**) Atl3-positive LCVs in the macrophages were quantified (n > 50 LCVs per condition and replicate). Shown are means and SD of three biological replicates. The percentage of Atl3-positive LCVs of infected and/or inhibitor-treated cells was compared to uninfected and untreated cells by two-way ANOVA with Tukey’s multiple comparisons test; ns, non-significant; ****, P<0.0001.

In another approach, we depleted IMPDH2 by RNA interference in A549 cells before infection with GFP-producing *L. pneumophila* wild-type or Δ*icmT* and assessed intracellular growth by relative GFP-fluorescence (**Fig. 5b**). Control oligonucleotides (“scrambled”) and siRNA targeting the ADP-ribosylation factor 1 (Arf1), were included as negative and positive controls, respectively, and the efficiency of IMPDH2 depletion was assessed by Western blot (**Fig S7c**). The increase of relative GFP-fluorescence within 24 h post-infection was about 40% or 50% reduced in wild-type *L. pneumophila*-infected host cells treated with siRNA targeting IMPDH2 or Arf1, respectively, as compared to host cells treated with “scrambled” siRNA. In summary, inhibition or depletion of IMPDH2 in host cells impairs intracellular growth of *L. pneumophila*, indicating that IMPDH2 promotes intracellular replication, which is in agreement with the notion that *L. pneumophila* exploits IMPDH2 activity during infection.

Since AnkX and its substrate IMPDH2 promote intracellular growth of *L. pneumophila*, we addressed the question whether these enzymes are involved in LCV formation. To this end, we infected RAW 264.7 macrophages with mCherry-producing wild-type, Δ*icmT* or Δ*ankX L. pneumophila* and lysed the infected cells 2 h post-infection, followed by immunostaining against the ER/LCV-marker atlastin-3 (Atl3) and analysis by fluorescence microscopy (**Fig. 5c**). LCVs formed in wild-type or Δ*ankX L. pneumophila-*infected macrophages similarly accumulated Atl3 2 h post-infection, indicating ER-acquisition, a hallmark of LCV formation (**Fig. 5d**). Likewise, treatment of the infected cells with the IMPDH2 inhibitors MPA or ribavirin did not alter Atl3-accumulation on the LCVs. Thus, neither the *Legionella* effector AnkX nor the host target IMPDH2 are required for LCV formation, yet both proteins bind the LCV. In the case of IMPDH2, the binding to LCVs depends on its oligomeric state, which is altered by AnkX-dependent phosphocholination, as observed *in vitro* and *in cellulo*.

### IMPDH2 is implicated in the modulation of host cell migration by *L. pneumophila*

IMPDH2 overexpression was shown to increase the migration of lung and colorectal cancer cells, while IMPDH2 depletion had opposite effects ^71,72^. Based on these findings, we investigated the involvement of AnkX in the migration of *L. pneumophila*-infected cells. To this end, we performed scratch assays with A549 cells infected with *L. pneumophila* wild-type, Δ*icmT*, Δ*ankX*, or Δ*ankX*/*ankX*, in absence or presence of the IMPDH2 inhibitors MPA (10 μM) or ribavirin (100 μM) (**Fig. 6a**). The IMPDH2 inhibitors were not toxic to host cells at the concentrations used (**Fig S7d**). 24 h after introducing a scratch to a monolayer of wild-type-infected cells only ca. 25% wound closure was observed (**Fig. 6b**). In contrast, for uninfected, Δ*icmT*-infected, or Δ*ankX*-infected monolayers, ca. 80-90% wound closure was seen, and providing the Δ*ankX* mutant strain with plasmid-borne *ankX* reduced wound closure. This suggests that AnkX largely contributes to the inhibition of cell migration caused by *L. pneumophila* infection. Treatment of the cells with IMPDH2 inhibitors decreased wound closure by ca. 15%, confirming the relevance of IMPDH2 for cell migration. Taken together, both the *Legionella* effector AnkX and its human target IMPDH2 affect cell migration, however in opposite ways.

**Figure 6.**
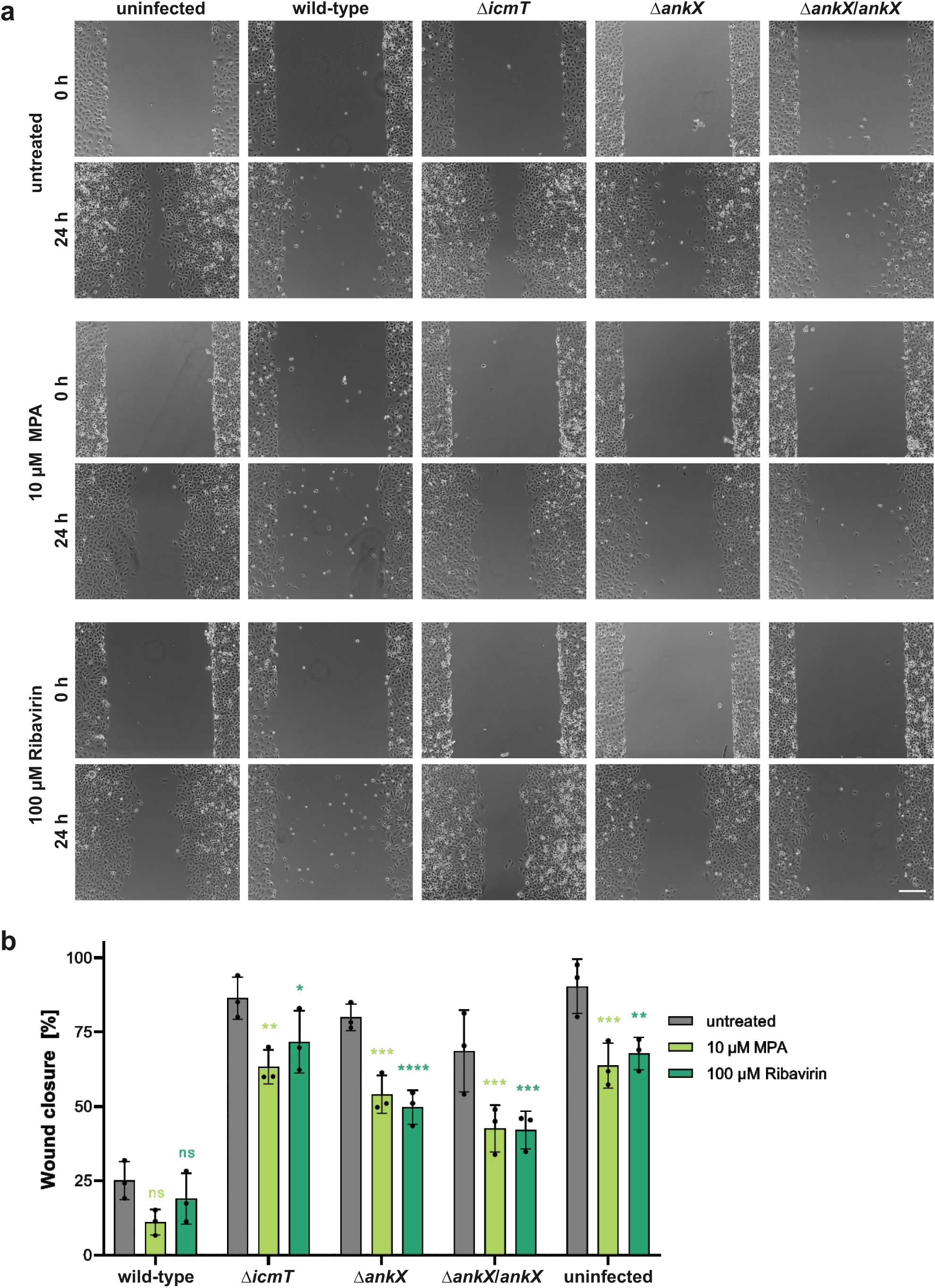
IMPDH2 promotes host cell migration. (**a**) Migration of A549 cells infected (MOI 20) with *L. pneumophila* wild-type, Δ*icmT*, Δ*ankX*, or Δ*ankX*/*ankX*, or treated with 10 µM MPA or 100 µM ribavirin was assessed by introducing a scratch to the cell monolayer 1 h after infection or treatment. The area of the scratch wound was determined by live imaging immediately after scratching and again after 24 h. (**b**) The wound closure was calculated for each treatment and compared to untreated and uninfected cells by a one-way ANOVA with Dunnett’s multiple comparisons test; *, P < 0.05; **, P < 0.01; ***, P < 0.001; ****, P < 0.0001; ns, non-significant. Plotted are individual values, means and SD of three independent experiments. Representative images of fixed samples (4% PFA) are shown. Scale bars, 200 µm.

## Discussion

In this study, we reveal that IMPDH2 promotes the intracellular growth of *L. pneumophila* (**Fig. 5**), and the effector AnkX phosphocholinates IMPDH2 (**Fig. 1**) at the residue Thr147 of the regulatory Bateman domain (**Fig. 3**). This covalent modification does not affect the activity of IMPDH2 *per se* (**Fig. S3a**) but modulates the nucleotide-dependent regulation of enzyme activity (**Figs. 1g, S3b**), oligomerization state (**Fig. 2**), cytoophidia dynamics (**Fig. 4**), and possibly subcellular localization (**Fig. 5**). Thus, *L. pneumophila* subverts host GTP metabolism by posttranslationally modifying a central enzyme of the GTP biosynthetic pathway.

AnkX has been initially characterized as a phosphocholine transferase targeting Rab1 and Rab35 ^23,25,69,73–75^. The phosphocholination of inactive GDP-bound Rab1 and Rab35 prevents their activation by guanine nucleotide exchange factors (GEFs) and binding to downstream interaction partners ^25^. Notably, AnkX phosphocholinates Rab1b on Ser76, whereas the structurally highly conserved Rab35 is modified at the homologous position on Thr76 ^23^. Accordingly, phosphocholination of IMPDH2 at Thr147 is consistent with the AnkX substrate scope. Intriguingly, the phosphocholinated threonine in human IMPDH2 (Thr147) is conserved and exposed in *D. discoideum* IMPDH (Thr153) (**Fig. S8**). This structural similarity suggests that AnkX not only phosphocholinates IMPDH at a threonine residue in mammalian cells but also in natural amoebae hosts.

Nevertheless, the phosphocholination of IMPDH2 and Rab1 shows some intriguing differences: (i) under the conditions used, IMPDH2 is phosphocholinated slower than Rab1 (**Fig. 1d**), and therefore, the phosphocholination kinetics are different, (ii) ATP (but not GTP) significantly and efficiently increases the phosphocholination of IMPDH2 but has no effect on the phosphocholination of Rab1 (**Fig. 1d**), and (iii) phosphocholinated IMPDH2, IMPDH2PC, is insensitive towards the dephosphocholinase Lem3, while phosphocholinated Rab1, Rab1_PC_, is readily dephosphocholinated by Lem3 (**Fig. S3c**). The latter finding is consistent with our previous work showing that Lem3 removes phosphocholine from serine, but not threonine residues ^26^. Accordingly, Lem3 acts on phosphocholinated Rab1b but not Rab35, and this selectivity is dictated by the modified residue rather than the Rab scaffold: exchanging the modified residue redirects Lem3 activity - Rab1b S76T is no longer dephosphocholinated, while Rab35 T76S becomes a Lem3 substrate ^26^. Thus, even if Lem3 were to bind IMPDH2, the T147 modification would likely remain resistant to its activity.

Based on these results, IMPDH2_PC_ is expected to experience a longer half-life compared to Rab1b_PC_ in *L. pneumophila*-infected cells. This differential reversibility may represent another layer of coordinated host-target manipulation by *Legionella* effectors. In this context, limiting the lifetime of Rab1b phosphocholination while permitting more persistent modification on threonine acceptors (Rab35 and IMPDH2) could be advantageous, because Rab1b is additionally AMPylated by the effector DrrA at Tyr77, directly adjacent to Ser76, and may therefore need to remain accessible for sequential modification and regulation during infection. Moreover, the differences in phosphocholination kinetics and half-lives of the two AnkX targets likely reflect the bi-phasic infection cycle of *L. pneumophila* ^76,77^. Initially, virulent and motile bacteria infect a host cell and instantly (within 1-2 h) form an LCV. LCV formation critically depends on the subversion of host cell vesicle trafficking, including the modulation of Rab1 and Rab35 implicated in (secretory) vesicle trafficking. Accordingly, AnkX contributes to a rapid modification of Rab GTPase targets, and due to the activity of the antagonist Lem3, phosphocholination of these targets is reversible. The initial LCV formation is followed by a phase switch: *L. pneumophila* adopts its replicative form and initiates intracellular growth. Depending mainly on the temperature, the bacteria grow intracellularly over the next 1-2 days until a subpopulation of virulent/motile bacteria lyses the LCV, exits from the host cell and spreads to new niches ^78^. At these later stages of infection, AnkX more slowly (and perhaps irreversibly) modifies IMPDH2, thus having a sustained impact on the GTP metabolism of the host cell, the biological implications of which are presently not clear.

The phosphocholination of IMPDH2 was promoted by ATP and slightly inhibited by GTP (**Fig. 1d**). Pre-incubation with ATP or the non-hydrolysable ATP analogue AMPPCP facilitated IMPDH2 phosphocholination and filament disassembly to similar levels (**Fig. 2cd**). In contrast, pre-incubation with GTP did not promote phosphocholination of IMPDH2, despite inducing filament formation (**Fig. 2cd**). Therefore, filamentation of IMPDH2 *per se* does not appear to facilitate AnkX-dependent phosphocholination. Rather, AnkX seems to preferably accept ATP-(or AMPPCP-) bound IMPDH2 as a substrate and perhaps specifically recognizes the extended octamer conformation.

The *L. pneumophila* effector protein AnkX phosphocholinates IMPDH2 at Thr147 (**Fig. 3**). Thr147 localizes to the regulatory Bateman domain of IMPDH2, which is not in direct proximity of the ATP and GTP binding sites but exposed to the surface of the enzyme (**Fig. S1a**). Thus, this site is readily accessible for post-translational modification(s). The Bateman domain is implicated in interactions with cellular partners such as ANKRD9 ^79^. Accordingly, the modification of IMPDH2 could perturb protein–protein interactions, as previously described for phosphocholinated Rab1 ^25^. Importantly, cytoophidia are thought to consist not solely of IMPDH2 but may serve as hubs where multiple enzymes cooperate within a synthetic pathway. Consequently, modification of IMPDH2 could broadly affect cellular metabolic organization, not only by homotypic filamentation but also by interacting with other proteins.

IMPDH2 cytoophidia were completely disassembled, upon *L. pneumophila* infection of A549 cells (**Fig. 4ab**), indicating that the process is of prime importance for an efficient bacterial infection. The disassembly of the cytoophidia was dependent on the Icm/Dot T4SS and the phosphocholinase AnkX. The almost complete disassembly within 2 h of cytoophidia composed of multiple densely packed IMPDH2 filaments indicates that phosphocholination represents a very efficient mechanism to modulate the structure and oligomeric organization of IMPDH2 and its cellular compartmentalisation.

Untreated IMPDH2 binds to LCVs, but the phosphocholination of ATP-treated IMPDH2 filaments barely increases binding of IMPDH2 to the pathogen compartment (**Figs 4c, S6c**). Still, the subcellular localization of IMPDH2 might be important for its cellular function, and *L. pneumophila* might benefit from redirecting IMPDH2 to LCVs or other cellular compartments. Accordingly, through the activity of AnkX, *L. pneumophila* might disassemble IMPDH2 filaments to increase the availability of non-filamentous IMPDH2 and to alter its subcellular localization. Overall, the stabilization of IMPDH2 in lower order oligomeric states might allow a more efficient re-distribution of the enzyme to cellular compartments and/or lock the enzyme in an infection-supportive conformation.

IMPDH2 promotes the intracellular growth of *L. pneumophila* (**Fig. 5ab**) but does not seem to affect the formation of the replication-permissive LCV (**Fig. 5c**). Neither the deletion of AnkX from the *L. pneumophila* genome nor the pharmacological inhibition of host IMPDH2 affected the accumulation of the ER-marker Atl3 on the LCV 2 h post-infection (**Fig. 5d**). Therefore, IMPDH2 (and GTP metabolism) are likely important for the intracellular bacteria after the LCV is established.

Other bacterial pathogens such as *Chlamydia trachomatis* ^80^ and *Brucella abortus* ^81^, as well as several viruses including yellow fever ^82^, chikungunya ^83^, and SARS-Cov-2 ^84^ viruses exploit the function of host cell IMPDH2 during infection. The molecular mechanism(s) underlying this requirement of IMPDH2 for intracellular growth of bacterial and viral pathogens are not known in detail. However, since IMPDH2 activity is crucial for GTP supply and replication of several intracellular pathogens, IMPDH2 inhibitors such as MPA or ribavirin are clinically applied to treat viral infections, cancer, autoimmune and inflammatory disorders ^47,85–89^.

Filament assembly of key metabolic enzymes is increasingly gaining attention as a mechanism of activity regulation. Recent work in structural and cell biology deepened our understanding of IMPDH2 filament formation and its effect on enzyme regulation and addressed the relevance of IMPDH2 cytoophidia in various cell types. However, the mechanisms controlling the reversible assembly of IMPDH2 into cytoophidia are incompletely understood and have not been investigated in the context of pathogen infection. In the present work, we showed that phosphocholination, a rare post-translational modification catalysed by the *L. pneumophila* effector AnkX, causes the depolymerisation of ATP-induced, linear IMPDH2 filaments *in vitro* (**Figs. 1-3**) as well as cytoophidia in *L. pneumophila*-infected cells, resulting in the subcellular redistribution of the enzyme (**Fig. 4**). Future studies will further address the (infection) biological implications of IMPDH2 and its post-translational covalent modifications.

## Materials and Methods

### Bacteria and cells

The bacterial strains used in this study are listed in **Table S1**. *L. pneumophila* strains were grown for three days at 37°C on charcoal yeast extract (CYE) agar plates buffered with N-(2-acetamido)-2-aminoethane sulfonic acid (ACES). 5-10 μg/mL chloramphenicol (Cam) was added if required. *L. pneumophila* over-night cultures were inoculated in ACES yeast extract (AYE) containing 5-10 μg/mL Cam at an OD_600_ of 0.1 for 21-22 h on a rotating wheel (37°C, 80 rpm) until a OD_600_ of ca. 5 was reached.

Human A549 lung epithelial cells (ATCC CCL-185, laboratory collection) and murine RAW 264.7 macrophages (ATCC TIB-71, laboratory collection) were cultured in RPMI 1640 medium supplemented with 10% fetal calf serum (FCS) and 2 mM L-glutamine (all from Gibco, Thermo Fisher), at 37°C and 5% CO_2_ in a humidified incubator.

### Bacterial growth assays

For assessment of bacterial growth in medium, *L. pneumophila* was grown in AYE for 21 h, diluted in AYE to an OD_600_ (optical density at 600 nm) of 0.1, and added at 200 µl per well to 96-well plates. Mycophenolic acid (MPA) was added to final concentrations of 0, 0.05, 0.5, 2.5, or 25 µM, and bacterial growth was measured by OD_600_ using a Synergy H1 Hybrid Reader (BioTek) while orbitally shaking at 37°C.

To assess intracellular replication of *L. pneumophila*, 2 × 10^4^ A549 cells per well were seeded into black 96-well plates (Corning) one day prior to the experiment. GFP-producing *L. pneumophila* wild-type, Δ*ankX*, Δ*lem3*, or Δ*icmT* harboring pNT28 or the complementation plasmids pLS284 or pLS285 were grown for 21 h in AYE and added to the cells at an MOI of 1. MPA was added to the infected cells to final concentrations of 0, 0.05, 0.5, 2.5, or 25 µM, and infection was synchronized by centrifugation (450 *g*, 10 min). The cells were incubated at 37°C and 5% CO_2_ in a humidified atmosphere for the duration of the experiment, and GFP fluorescence was measured as relative fluorescent units (RFUs) every 24 h using a Cytation 5 Cell Imaging Multi-Mode Reader (BioTek).

### Molecular cloning and construction of *L. pneumophila* mutant strains

Plasmids and oligonucleotides used in this study are listed in **Table S1** and **Table S2**, respectively. Plasmids, genomic DNA and PCR products were purified using the NucleoSpin Plasmid Mini kit for plasmid DNA (Macherey-Nagel) or the NucleoSpin Gel and PCR Clean-up Kit (Macherey-Nagel). PCRs were performed using Phusion High-Fidelity DNA polymerase (Thermo Fisher), and plasmid backbones were digested using restriction enzymes from New England Biolabs (NEB) or Thermo Fisher according to the manufacturer’s protocols. Insertion of PCR products into backbones was performed by Gibson assembly using the NEBuilder HiFi DNA assembly kit (NEB). Correct PCR amplification was verified by sequencing.

Chromosomal deletion of *ankX* and *lem3* were performed as previously described ^90^. In short, 1,000 bp upstream and downstream fragments of *ankX* and *lem3* were PCR amplified using genomic DNA of *L. pneumophila* strain Philadelphia-1 as a template. Primer pairs oLS027/oLS028 and oLS029/oLS030 were used for the respective *ankX* flanking regions and primer pairs oLS069/oLS051 and oLS052/oLS085 for the *lem3* flanking regions. The up-and downstream fragments together with a Kan^R^ resistance cassette, from pUC4K, were directly inserted into the pLAW344 suicide plasmid by a four-way ligation. For the *ankX* deletion plasmid, pLS030, the fragments were ligated into pLAW344 with SalI and PstI for the Kan^R^ cassette. For the *lem3* deletion plasmid, pLS098, the fragments were ligated using BamHI and SalI for the Kan^R^ cassette. Plasmids were confirmed by sequencing and transformed by electroporation into *L. pneumophila* JR32. Mutant clones were isolated by simultaneous selection for Kan^R^ and Suc^R^, and correct insertion of the Kan^R^ resistance cassette was confirmed by PCR screening and sequencing.

Plasmids pLS284 and pLS285 for complementation of the Δ*ankX* and Δ*lem3* deletion mutants were constructed by PCR amplification of the respective gene together with its upstream 600 bp promotor region using the primer pairs oLS441 and oLS442, or oLS443 and oLS444, respectively. PCR products were cloned into BamHI-digested pNT31 backbone by Gibson assembly.

The IMPDH2 construct, containing the full length IMPDH2 (UniProt entry P12268), was cloned into a pMAL vector (New England Biolabs) in which the N-terminal His_6_-and MBP (maltose-binding protein)-Tag is separated from IMPDH2 by a TEV (tobacco etch virus) protease cleavage site. Plasmid pLS281 for expression of IMPDH2 in *E. coli* was constructed by amplification of human *impdh2* from an *impdh2* ORF cDNA clone in a cloning vector (LubioScience) using the primers oLS438 and oLS439. Plasmid pRH003 was modified by quick change, as previously described ^91^, to introduce a KpnI restriction site (primers oRH082 and oRH083) and to remove the thrombin cleavage site (primers oRH086 and oRH087). The resulting backbone was digested with KpnI and SalI and assembled with the amplified *impdh2* gene by Gibson assembly. Plasmids pLS282, pPS016, and pPS017 were constructed by quick change using pLS281 as a template. Primers oLS419 and oLS420 were used to introduce a T147A (ACA to GCA) mutation (pPS016), primers oLS421 and oLS422 to introduce a S160A (TCC to GCC) mutation (pPS017) and both primer pairs to mutate both sites (pLS282).

Plasmid pPS014 for expression of AnkX_1-800___H229A_ was constructed by quick change using the primers oPS049 and oPS050 to introduce a CAC to GCC mutation and pSF421 as a template. The His_10_-GFP-tagged AnkX constructs, containing the amino acids 1-800, was previously described in ^75^. The His_10_-GFP-tagged Lem3 construct, containing the amino acids 21-486, as well as the Rab1b construct, containing the amino acids 3-174, was previously described in ^26^.

### Pulldown of biotinylated proteins and in-solution digestion

Eukaryotic proteins phosphocholinated by AnkX were identified by covalent biotinylation using a CDP-choline-biotin conjugate (**Fig. 1a**), which was synthesized as published ^65^ via a five-step sequence starting from 1-bromo-carboxybenzylamine-PEG₄. For each phosphocholination experiment, 1 mg of total protein lysate was transferred to a microcentrifuge tube and adjusted to a total volume of 250 µL with lysis buffer (50 mM PIPES, 50 mM NaCl, 5 mM MgCl₂, 5 mM EGTA, 0.1% Tween-20, 0.1% NP-40, 0.1% Triton X-100, pH 7.4). The CDP-choline-biotin probe (final concentration 100 µM) and AnkX (final concentration 1 µM) were added, followed by incubation for 24 h at 4 °C under gentle agitation. Control samples were prepared in parallel without AnkX.

Biotinylated proteins were identified by Western blot. To this end, samples from the labeling reaction were taken after distinct time points and quenched in Laemmli buffer at 95°C. Subsequently, samples were separated by SDS-PAGE and transferred onto a nitrocellulose membrane using a Trans-Blot® TurboTM Blotting system (Bio-Rad). The membrane was blocked in Tris buffered saline, containing 0.1% Tween20 (TBS-T) and 1xRoti®-Block (Roth) for 0.5 h. Subsequently, a streptavidin-fluorescein conjugate (Thermo Fisher) was pipetted into the blocking solution (1:1000) and incubated overnight at 4°C. The membrane was washed three times with TBS-T for 10 minutes and fluorescein-signals were detected on a fluorescence imaging system (Odyssey, LI–COR Biosciences).

To identify target proteins by Proteins were precipitated by adding ice-cold acetone (approx. 1 mL) and incubating for 30 min at-20 °C. The precipitate was collected by centrifugation (13,000 rpm, 10 min, 4°C), washed three times with ice-cold methanol, and resuspended in 500 µL PBS containing 0.2% SDS by brief sonication. Streptavidin magnetic beads (New England Biolabs, #S1420S) were washed twice with PBS, and 250 µL of bead suspension was added to each sample. The mixtures were incubated for 1 h at room temperature (RT) under rotary agitation to capture biotinylated proteins. Beads were then washed sequentially with 3 × 1 mL PBS containing 0.2% SDS, 2 × 1 mL 6 M urea, and 8 × 1 mL PBS to remove nonspecific binders.

For on-bead digestion, beads were resuspended in 100 µL elution buffer E1 (50 mM ammonium bicarbonate, 2.2 M urea, 1 mM DTT, 5 µg trypsin) and incubated for 1 h at RT (thermomixer, 350 rpm). The supernatant was transferred to a fresh tube, and the beads were washed with 100 µL elution buffer E2 (50 mM ammonium bicarbonate, 2.2 M urea, 5 mM chloroacetamide) for 2 min at RT. The E2 wash was combined with the E1 fraction, and digestion was continued overnight at 37 °C in a thermomixer (350 rpm). The reaction was quenched by adding 2 µL concentrated trifluoroacetic acid (TFA) per sample.

Peptides were desalted using C18 StageTips prepared from 3M Empore C18 extraction disks (47 mm, cat. #2215). Each tip was activated with 100 µL methanol and equilibrated twice with 100 µL buffer A (0.1% formic acid in water). 100 µL pull down sample were loaded onto the tips, incubated for 1 min, and centrifuged (5,000 rpm). The tips were washed once with 100 µL buffer A (centrifuged at 5,000 rpm) and peptides were eluted twice with 20 µL buffer B (80% acetonitrile, 20% water, 0.1% formic acid) by centrifugation (5,000 rpm). Eluates were dried in a SpeedVac at RT and stored at-20 °C until LC–MS/MS analysis.

### NanoHPLC-MS/MS analysis of tryptic peptides

For protein identification and quantification, the tryptic peptides were separated and analyzed by nano-HPLC-MS/MS. The analyses were perfomed on an Ultimate 3000 nano-HPLC coupled to an Orbitrap Q-Exactive mass spectrometer (ThermoFisherScientific, Germany). All solvents were LC-MS grade.

Dried tryptic peptides were dissolved in 20 µl 0.1 % TFA. 3 µl of these samples were injected and enriched onto a C18 PepMap 100 column (3 µm, 100 Å, 300 µm ID * 5 mm, Dionex, Idstein, Germany) using 0.1 % TFA and a flow rate of 30 µl/min for 5 min. Peptides were separated on a C18 PepMap 100 column (3 µm, 100 Å, 75 µm ID * 250 mm) using a linear gradient starting with 95% solvent A / 5% solvent B and increasing to 30.0% solvent B in 90 min with a flow rate of 300 nl/min (solvent A: water containing 0.1 % formic acid; solvent B: acetonitrile containing 0.1 % formic acid). The nano-HPLC was online coupled to the Orbitrap Q-Exactive mass spectrometer using a standard coated Pico Tip (ID 20 µm, Tip-ID 10 µM, New Objective, Woburn, MA, USA). Precursor scans were carried out with a resolution of 70,000. MS/MS data of the 10 most intense and at least two-fold charged ions were recorded with a resolution of 17500.

Data analysis was performed using MaxQuant (v 2.6.7.0) ^92^, including the Andromeda search algorithm and searching the human and *L. pneumophila* reference proteomes of the Uniprot database. The search was performed for full enzymatic trypsin cleavages allowing two miscleavages. For protein modifications, carbamidomethylation was chosen as fixed and oxidation of methionine and acetylation of the N-terminus as variable modifications. The mass accuracy for full mass spectra was set to 20 ppm (first search) and 4.5 ppm (second search), respectively and for MS/MS spectra to 20 ppm. The false discovery rates for peptide and protein identification were set to 1%. The option “Match-between-runs” was activated. Only proteins for which at least two peptides were quantified were chosen for further validation. Relative quantification of proteins was performed by using the label-free quantification algorithm implemented in MaxQuant.

Further statistical data analysis was performed using Perseus (v.2.0.11.0) ^93^. Normalized Label-free quantification (LFQ) intensities were log-transformed (log2) and grouped (control vs. sample). Proteins had to be quantified in all three replicates in at least one of the groups to be retained for further analysis. Missing values were imputed by normally distributed small values (width 0.3, down shift 1.3) and a two-sided Student’s t-Test (permutation-based corrected, s0=0) was performed. All proteins with log2 fold change (FC) greater than 3 and p-values less than 0.01 were considered as statistically significant.

### Protein production and purification

For negative stain transmission electron microscopy (TEM), phosphocholination kinetics measurements, and LCV binding experiments, human IMPDH2 and its mutants IMPDH2_T147A_, IMPDH2_S160A_, and IMPDH2_T147A_S160A_ were produced in *E. coli* BL21-CodonPlus (DE3)-RIL transformed by heat-shock transformation with plasmids pLS281, pPS016, pPS017, or pLS282, respectively. A preculture starting with an OD_600_ of 0.1 was grown for 3 h at 37°C and 160 rpm in lysogeny broth (LB) containing 50 μg/ml Kan and 30 μg/ml Cam. Main cultures (600 ml) were inoculated at a starting OD_600_ of 0.075 in LB, supplemented with Kan and Cam, and grown at 37°C and 95 rpm until OD_600_ 0.6. Protein production was induced with 1 mM isopropyl-β-d-thiogalactoside (IPTG; Carl Roth) and performed overnight at 21°C and 85 rpm.

To produce the bacterial effector protein AnkX_1-800_ and its catalytically inactive mutant AnkX_1-800___H229A_, *E. coli* BL21(DE3) were transformed with plasmids pSF421 or pPS014, respectively. Precultures were grown overnight at 37°C and 160 rpm in LB containing 100 μg/mL Ampicillin (Amp). Main cultures (600 ml) with a starting OD_600_ of 0.03 were grown at 37°C and 95 rpm until OD_600_ 0.1, then the temperature was reduced to 20°C. After reaching OD_600_ 0.6, overnight protein production was induced with 0.5 mM IPTG. Bacterial cultures were harvested by centrifugation (6,000 *g*, 20 min, 4°C), pooled by resuspension in LB, pelleted (600 *g*, 4°C, 20 min) and snap-frozen in liquid nitrogen for storage at-80°C.

For purification of IMPDH2 and its mutants, pellets were resuspended in lysis buffer A (50 mM KH_2_PO_4_, 300 mM KCl, 800 mM urea, pH 8) ^67^ supplemented with protease inhibitors (Pierce Protease Inhibitor Mini Tablets, EDTA-free; Thermo Fisher) and DNAse (Sigma Aldrich). AnkX_1-800_-and AnkX_1-800___H229A_-producing bacteria were resuspended in lysis buffer B (50 mM Tris, 500 mM KCl, 10% glycerol, 2 mM β-mercaptoethanol, pH 8) supplemented with 1 mM PMSF, 3 mM MgSO_4_, and DNAse (Sigma Aldrich). Bacterial cells were lysed by high-pressure homogenization (Microfluidizer; Microfluidics, 21,000 psi), lysates were centrifuged (16,000 *g*, 4°C, 30 min), supernatants were supplemented with 10 mM imidazole (Sigma Aldrich) and passed through Ni^2+^-NTA affinity chromatography columns. Columns were washed with lysis buffer A (for IMPDH2 and mutants) or lysis buffer B (AnkX_1-800_-and AnkX_1-800___H229A_) supplemented with 50 mM imidazole, and proteins were eluted with elution buffer A (50 mM KH_2_PO_4_, 300 mM KCl, 800 mM urea, 500 mM imidazole, pH 8), for IMPDH2 and its mutants, or with elution buffer B (50 mM Tris, 500 mM KCl, 10% glycerol, 2 mM β-mercaptoethanol, 300 mM imidazole, pH 8) for AnkX_1-800_ and AnkX_1-800___H229A_. His-tagged IMPDH2 and mutants, as well as His-GFP-tagged AnkX_1-800_ and AnkX_1-800___H229A_ were concentrated using Amicon Ultra filters with 30 or 100 kDa molecular weight cut off (MWCO), and buffer was exchanged for lysis buffer A or lysis buffer B, respectively, using PD-10 columns (Cytiva). Proteins were incubated overnight at 4°C with 3C protease (produced in-house, 10 μg/mL) to cleave off the His-tag, or with TEV protease (Sigma, 20 μg/mL) to cleave off the His-GFP-tag and passed through Ni^2+^-NTA affinity chromatography columns to remove cleaved off tags.

For aSEC, the proteins were concentrated using 30 kDa MWCO (IMPDH2 and mutants) or 50 kDa MWCO (AnkX_1-800_ and AnkX_1-800___H229A_) Amicon Ultra filters, and loaded onto a Superose 6 Increase 10/300 GL (IMPDH2 and mutants) or a Superdex 200 Increase 10/300 GL (AnkX_1-800_ and AnkX_1-800___H229A_) column (Cytiva) equilibrated with gel filtration buffer A (20 mM HEPES, 100 mM KCl, 1 mM DTT, pH 8) or gel filtration buffer B (50 mM Tris, 500 mM KCl, 2 mM β-mercaptoethanol, pH 8), respectively. Peak elution fractions were pooled and supplemented with 10% glycerol, and protein aliquots were snap-frozen in liquid nitrogen for storage at-80°C. Protein concentrations were determined by OD_280_ measurements (NanoDrop 2000) and calculated using the respective extinction coefficient (https://www.expasy.ch/tools/protparam.html). Protein purity and concentrations were routinely confirmed by SDS-PAGE.

### Preparative IMPDH2 production and purification

AnkX and IMPDH2 constructs were produced in *E. coli* BL21-CodonPlus (DE3)-RIL cells. At first, cells were precultured in LB for 4 h at 37°C and 200 rpm (Infors HT shakers) before main cultures were inoculated with a starting OD_600_ of 0.05. Main cultures of 1 L volume were grown at 37 °C and 180 rpm until OD_600_ 0.6 to 0.8, and protein production was induced by adding 0.5 mM IPTG. Protein production was performed at 21°C for 16 to 18 h. Afterwards cells were harvested by centrifugation (4,000 *g*, 20 min) and directly solved in the respective buffer A for purification: buffer A_IMPDH2 (50 mM HEPES-NaOH pH 8, 500 mM KCl, 5% Glycerol, 2 mM TCEP), buffer A_AnkX (50 mM HEPES-NaOH pH 8, 500 mM NaCl, 5% Glycerol, 2 mM TCEP), the respective buffer B was additionally supplemented with 500 mM imidazole. Prior to cell lysis, a spatula tip of DNAse I (Sigma-Aldrich) was added to the bacteria resuspended in buffer A. Cell lysis was performed using a French press system (Constant Cell Disruption Systems) at 1.8 kbar, and endogenous protease activity was inhibited by adding 1 mM phenylmethylsulfonyl fluoride (PMSF). Cell lysates were cleared by centrifugation (21,000 *g*, 45 min) and supplemented with 25 mM imidazole.

Chromatography steps were performed running 5 mL Nuvia IMAC column (BioRad), 5 mL MBPTrap HP column (GE Healthcare Life Sciences) and HiLoad 16/600 Superdex columns (200 pg) (Cytiva) on an NGC medium-pressure liquid chromatography (LC) system (BioRad). Initial separation of endogenous *E. coli* proteins and the protein of interest was achieved performing immobilized metal affinity chromatography (IMAC) with a washing step of 30 mM (IMPDH2) or 40 mM (AnkX) imidazole for 40 column volumes and protein elution at 125-150 mM imidazole. Dialysis was performed over night at 4 °C against respective dialysis buffer (IMPDH2: 20 mM HEPES-NaOH pH 8, 300 mM KCl, 5% glycerol, 0.5 mM TCEP; AnkX: 20 mM HEPES-NaOH pH 8, 100 mM NaCl, 5% glycerol, 0.5 mM TCEP) in presence (IMPDH2) or absence (AnkX) of TEV protease. AnkX was concentrated to desired concentration using Amicon Ultra 15 mL centrifugal filters (Merck Millipore) directly after dialysis, while IMPDH2 was further purified. The cleaved His_6_-MBP-Tag was separated from IMPDH2 using an MBP-Trap column in combination with the IMAC to also separate the TEV protease. In a last step, IMPDH2 was transferred to storage buffer (20 mM HEPES-NaOH pH 7.5, 300 mM KCl, 5% glycerol, 0.5 mM TCEP) by aSEC and concentrated as described for AnkX. Proteins were stored at-80°C.

### Mass spectrometry of intact proteins

For intact mass spectrometry (iMS), samples were desalted using C18 ZipTips (Millipore, USA) and analyzed in MeOH:2-PrOH:0.2% FA (30:20:50). The solution was infused through a fused silica capillary (ID75 µm) at a flow rate of 1 µL/min and sprayed through a Pico Tip (ID30 µm; CoAnn Technologies, Richland WA, USA). Nano ESI-MS analysis of the sample was performed on a Synapt G2_Si mass spectrometer, and the data were recorded with the MassLynx 4.2 Software (both Waters, UK). Mass spectra were acquired in the positive-ion mode by scanning an m/z range from 400 to 5,000 Da with a scan duration of 1 s and an interscan delay of 0.1 s. The spray voltage was set to 3 kV, the cone voltage up to 100, and source temperature 100°C. The recorded m/z data were then deconvoluted into mass spectra by applying the maximum entropy algorithm MaxEnt1 (MaxLynx) with a resolution of the output mass 0.5 Da/channel and Uniform Gaussian Damage Model at the half height of 0.5 Da.

Alternatively, iMS was performed on a maXis II ETD ESI MS (Bruker Daltonics) device coupled to an Elute UHPLC (Bruker Daltonics). Purified proteins were desalted prior to MS analysis by injection onto a ProSwift™ RP-4H 1×50 mm column (Thermo Fisher) running at 0.3 mL min^-1^ and subsequent elution in an acetonitrile-gradient (5-90%) over a period of 2 min and 20 sec. IMPDH2 was injected at a final amount of 0.6 μg. Data were analysed using DataAnalysis (Version 5.1, Bruker Daltonics).

### Identification of the phosphocholination site by tandem mass spectrometry

#### Protein extraction and tryptic digestion

20 µg of the samples were dissolved to a concentration of 70% acetonitrile (ACN). 2 µL carboxylate modified magnetic beads (GE Healthcare Sera-Mag™, Chicago, USA) at 1:1 (hydrophilic/hydrophobic) in methanol were added following the SP3-protocol workflow ^94^. Samples were shaken (1400 rpm, 18 min, RT). The beads were bound to a magnetic rack, and the supernatant was removed. Magnetic beads were washed twice with 100% ACN, twice with 70% Ethanol, and resuspended in 50 mM ammonium bicarbonate. Subsequently, disulfide bonds were reduced with 10 mM dithiothreitol for 30 min, alkylated in presence of 20 mM iodoacetamide for 30 min in the dark and digested with trypsin (sequencing grade, Promega) at 1:100 (enzyme:protein) at 37 °C overnight while shaking at 1400 rpm. For binding of tryptic peptides, the beads were dissolved to 95% ACN and shaken (1400 rpm, 10 min, RT). Tubes were placed on the magnetic rack, the supernatant was removed, and the beads were washed twice with 100% ACN. Elution was performed with 2% DMSO in 1% formic acid (FA). The supernatant was dried in a vacuum centrifuge and stored at-20°C until further use.

#### LC settings

Prior to LC-MS/MS analysis, samples were dissolved in 0.1% FA to a final concentration of 1 µg/µL. For liquid-chromatography-coupled tandem mass spectrometry (LC-MS/MS) measurements, 0.25 µg tryptic peptides were injected for individual samples.

Measurements were performed on a quadrupole-ion-trap-orbitrap MS (Orbitrap Fusion, Thermo Fisher) coupled to a nano-UPLC (Dionex Ultimate 3000 UPLC system, Thermo Fisher). Chromatographic separation of peptides was achieved with a two-buffer system (buffer A: 0.1 % FA in water, buffer B: 0.1 % FA in ACN). Attached to the UPLC was a peptide trap (100 μm × 200 mm, 100 Å pore size, 5 μm particle size, C18, Thermo Fisher) for online desalting and purification followed by a 25 cm C18 reversed-phase column (75 μm × 250 mm, 130 Å pore size, 1.7 μm particle size, Peptide BEH C18, Waters). Peptides were separated using an 80-min gradient with linearly increasing ACN concentration from 2 % to 30 % ACN in 60 min. Eluting peptides were ionized using a nano-electrospray ionization source (nano-ESI) with a spray voltage of 1800, transferred into the MS and analyzed in data dependent acquisition (DDA) mode.

CID/HCD Measurement (MS^3^): For each MS1 scan, ions were accumulated for a maximum of 120 msec or until a charge density of 2 × 10^5^ ions (AGC Target) was reached. Fourier-transformation based mass analysis of the data from the orbitrap mass analyzer was performed covering a mass range of 400-1300 m/z with a resolution of 120,000 at m/z = 200. Peptides with charge states between 2+ - 5+ above an intensity threshold of 10,000 were isolated within a 2.5 m/z isolation window in Top Speed mode for 3 sec from each precursor scan and fragmented with a normalized collision energy of 30 % using CID. MS2 scanning was performed, using an ion trap mass analyzer at a rapid scan rate, with a start mass range of 120 m/z, and accumulated for 60 msec or to an AGC target of 1×10^4^. Peptides already fragmented were excluded for 30 sec. Targeted mass fragmentation using 30% HCD was triggered for m/z 187.07 within an isolation window of 2 m/z. MS3 scanning was performed on the orbitrap at a mass resolution of 30,000 over a scan range of 50-200 m/z and accumulated for 54 msec or to an AGC target of 10,000 with 1 scan.

ETD Measurement (MS^2^): For each MS1 scan, ions were accumulated for a maximum of 120 msec or until a charge density of 2 × 10^5^ ions (AGC Target) was reached. Fourier-transformation based mass analysis of the data from the orbitrap mass analyzer was performed covering a mass range of 200-1300 m/z with a resolution of 120,000 at m/z = 200. Peptides with charge states between 2+ - 5+ above an intensity threshold of 10,000 were isolated within a 1.6 m/z isolation window in Top Speed mode for 3 sec from each precursor scan and fragmented with ETD for a reaction time of 50 msec and a maximum of 200 msec ETD reagent injection time. MS2 scanning was performed, using an ion trap mass analyzer at a rapid scan rate, with a start mass range of 120 m/z, and accumulated for 60 msec or to an AGC target of 1×10^4^. Peptides already fragmented were excluded for 30 sec.

#### Raw data processing

LC-MS/MS from ETD measurements were searched with the Sequest algorithm integrated in the Proteome Discoverer software (v 2.4.1.15), Thermo Fisher) against the (modified) IMPDH2 protein sequence and a contaminant database. Carbamidomethylation was set as fixed modification for cysteine residues, and the oxidation of methionine, pyro-glutamate formation at glutamine residues at the peptide N-terminus, as well as acetylation of the protein N-terminus and phosphocholination (PC) of His, Ser, and Thr (+165.0554 Da) were allowed as variable modifications. A maximum number of 2 missing tryptic cleavages was set. Peptides between 6 and 144 amino acids were considered. A strict cutoff (FDR < 0.01) was set for peptide and protein identification.

For CID/HCD analysis, spectra were analyzed manually in QualBrowser within the Thermo Xcalibur software (v4.2.47, Thermo Fisher). For verification of ETD spectra, diagnostic ions generated by CID and HCD fragmentation of the phosphocholination were searched. Specifically, MS1: m/z 184 and MS2: m/z 86, 125, and 148, as published previously, were examined ^23^. Precursor m/z and retention times were compared to ETD measurements for confirmation.

### Phosphocholination and dephosphocholination of IMPDH2/IMPDH2_PC_

Preparative phosphocholination of IMPDH2 was performed in modification buffer (20 mM HEPES-NaOH pH 7.5, 20 mM NaCl, 1 mM MgCl_2_, 100 μM ATP, 5 mM CDP-choline, 5% glycerol, 2 mM TCEP) using purified His_10_-GFP-AnkX. IMPDH2 was diluted in modification buffer to a final concentration of 5 μM and preincubated at 19°C for 10 min before His_10_-GFP-AnkX was added at a final concentration of 5 μM. Phosphocholination of IMPDH2 was monitored using iMS over a period of 4-12 h. At a modification rate of 85% or higher, the reaction was diluted in buffer A_IMPDH2, supplemented with 40 mM imidazole, and IMPDH2 was separated from AnkX using IMAC. Finally, aSEC was performed and fractions containing IMPDH2 modified to 90% or higher were collected, concentrated to the desired concentration and stored at-80°C.

Analytical dephosphocholination of IMPDH2_PC_ by Lem3_21-486_ was tested in presence and absence of 100 μM ATP and/or GTP. GDP-loaded Rab1b_S76(PC)_ was used as a positive control for Lem3_21-486_ catalytic activity ^26^. Phosphocholinated substrates were co-incubated with Lem3_21-486_ for 24 h at 18°C and subsequently measured using iMS. As an internal reference the three biological replicates of IMPDH2 were also incubated without Lem3_21-486_ under the same conditions and measured using iMS. Intensities of IMPDH2 and IMPDH2_PC_ were compared within each sample to calculate the modification rate.

### IMPDH2 activity assay

The IMPDH2 activity (NAD⁺-dependent conversion of IMP to XMP) was assessed by measuring the NADH absorbance at 340 nm over time using a TECAN SPARK plate reader (Tecan). IMPDH2 was diluted to a final concentration of 2 μM in reaction buffer (HEPES-NaOH pH 7.5, 100 mM KCl, 10 μM MgCl_2_, 5% glycerol, 100 μM TCEP) and preincubated with 50 μM of IMP and varying concentrations of ATP or GTP (25, 5, 1, 0.2 mM) for 15 min at 18°C in Immuno Clear Standard Modules (Thermo Fisher). The reaction was started by the addition of 50 μM of NAD^+^. Resulting curves were used to determine initial reaction velocities and converted to the NADH production rate over time. P-values were calculated in an unpaired t test using the online T test calculator from GraphPad.

### Correlation of filament disassembly with IMPDH2 phosphocholination

#### Negative stain electron microscopy

For TEM of protein samples, purified IMPDH2 was diluted to 5 μM in phosphocholination buffer (20 mM Tris pH 7.0, 100 mM NaCl, 1 mM MgCl_2_, 1 mM DTT). Filament formation was induced by addition of 1 mM ATP (ATP magnesium salt, Sigma Aldrich) for 1 h at RT. To assess filament formation and phosphocholination in presence of different nucleotides, ATP was substituted with 1 mM IMP (inosine 5′-monophosphate disodium salt hydrate, Sigma Aldrich), 1 mM GTP (GTP sodium salt hydrate, Sigma Aldrich), or 1 mM AMPPCP (adenosine-5’-[(β,γ)-methyleno]triphosphate sodium salt, Jena Bioscience). When using AMPPCP, samples were additionally supplemented with 1 mM MgCl_2_. After incubation with nucleotides, 0.5 μM purified AnkX and 100 μM CDP-choline (CDP-choline sodium salt dihydrate, Sigma Aldrich) were added, and samples were further incubated for 2 h at RT. For negative controls, the catalytically inactive mutant AnkX_H229A_ was added instead, or either AnkX or CDP-choline, or both were omitted. For comparison of the point mutants IMPDH2_T147A_, IMPDH2_S160A_, and IMPDH2_T147A_S160A_ to wild-type IMPDH2, the proteins were diluted to 5 μM in phosphocholination buffer, incubated for 1 h at RT with 1 mM ATP for induction of filament formation, then for additional 2 h in the presence of 0.167 μM AnkX and 100 μM CDP-choline.

Negative staining was performed by placing samples, diluted 1:5 in phosphocholination buffer, onto glow-discharged (1 min) carbon film coated copper grids, 300 mesh (Science Services). After 1 min incubation, the sample was removed from the grid with a filter paper, and grids were immediately washed with 1% uranyl acetate, blotted, and stained with 1% uranyl acetate for 1 min. Dried grids were imaged by TEM with a FEI Tecnai Spirit at 120 kV.

#### Mass spectrometry

Protein samples subjected to intact mass spectrometry (iMS) were prepared analogous to the samples analyzed by TEM. Mass spectrometric analysis was performed by the Functional Genomics Center Zurich (FGCZ) of University of Zurich and ETH Zurich. Samples were desalted using C18 ZipTips (Millipore, USA) and analyzed in MeOH:2-PrOH:0.2% FA (30:20:50). The solution was infused through a fused silica capillary (ID75µm) at a flow rate of 1 µL min^-1^ and sprayed through a Pico Tip (ID30µm) obtained from CoAnn Technologies (Richland WA, USA). Nano ESI-MS analysis of the sample was performed on a Synapt G2_Si mass spectrometer, and the data were recorded with the MassLynx 4.2 Software (both Waters, UK). Mass spectra were acquired in the positive-ion mode by scanning an m/z range from 400 to 5,000 da with a scan duration of 1 sec and an interscan delay of 0.1 sec. The spray voltage was set to 3kV, the cone voltage to 50, and source temperature 100 °C. The recorded m/z data were then deconvoluted into mass spectra by applying the maximum entropy algorithm MaxEnt1 (MaxLynx) with a resolution of the output mass of 0.5 Da/channel and Uniform Gaussian Damage Model at the half height of 0.5 Da.

### Analytical size exclusion chromatography

For analytical size exclusion chromatography (aSEC), 100 μg of purified protein and 50 μM Vitamin B12 used as a standard in a final volume of 90 μl were injected onto a Superdex 200 10/300 column (Cytiva) connected to a Shimadzu HPLC System. The aSEC was performed in 20 mM HEPES-NaOH pH 7.5, 300 mM KCl, 5% glycerol, 0.5 mM TCEP at a flow rate of 0.4 mL min^-1^, and proteins were detected dually at 280 nm and 340 nm. If indicated, proteins were preincubated with 1 mM ATP for 60 min at 8°C prior to injection. The modification rate of IMPDH2 as indicated for each sample was created by mixing almost fully modified IMPDH2 and nonmodified IMPDH2 and assessed by iMS for each individual sample. All runs were compared to a standard (1,35 - 600 kDa, BioRad).

### Dynamic light scattering

IMPDH2 was diluted in modification buffer to a final concentration of 10 μM, and nucleotides were added at a final concentration of 100 μM as indicated. 10 μl sample were loaded into a glass capillary (Prometheus NT.48 Series nanoDSF Grade Standard Capillaries, Technologies GmbH) and measured in a Prometheus Panta device from NanoTemper with 100% LED and Laser power at 15°C. Per sample, 10 consecutive measurements were performed (technical replicates, final n = 30 per biological sample). Data were analysed using the Panta Analysis Software that marked outliers automatically and excluded them from further analysis steps.

### Immunofluorescence microscopy

#### Cytoophidia formation in infected cells

1 × 10^5^ A549 cells per well were seeded into 24-well plates (Corning) on top of sterilized 12 mm cover slips (Carl Roth) one day prior to the experiment. The following day, the cells were treated with 500 nM MPA and simultaneously infected (MOI 100) with GFP-producing (pNT28 or pLS284) *L. pneumophila* wild-type, Δ*icmT*, Δ*ankX*, or Δ*ankX*/*ankX* grown in AYE for 21 h, or left uninfected. Infections were synchronized by centrifugation (450 *g*, 10 min), and cells were incubated for 50 min at 37°C and 5% CO_2_ in a humidified atmosphere. After 1 h, cells were washed twice with RMPI 1640 medium and further incubated for 1 h in the presence of 500 nM MPA. At 2 h post-infection, cells were washed with DPBS, fixed with 4% PFA (20 min 37°C), and permeabilized with 0.1% Triton X-100 in DPBS (30 min, RT). Following cell permeabilization, coverslips were blocked with 1% BSA in DPBS (1 h, RT), removed from the wells and incubated overnight at 4°C with rabbit anti-IMPDH2 antibody (Abcam: ab131158) diluted 1:250 in 1% BSA in DPBS. After primary antibody staining, coverslips were washed in DPBS and incubated for 1 h at RT with anti-rabbit Alexa 594 antibody (Invitrogen: A-21442) diluted 1:250 in 1% BSA in DPBS. After secondary antibody staining, coverslips were again washed in DPBS and mounted onto microscope slides (Carl Roth) with ProLong Diamond Antifade Mountant with DAPI (Thermo Fisher). Samples were left to dry overnight at RT and imaged using a Leica SP8 DMi8 CS with AFC laser scanning microscope, an HC PL APO CS2 63×/1.4 oil objective, and Leica LAS X software (Leica). For each condition, 16 tile-scans were acquired as z-stacks with 1× zoom, 0.28 µm z-step increments, bi-directional laser scan and a scanning speed of 500 Hz. Tile-scans were merged in the Leica LAS X software, and multiple intensity projections were computed in ImageJ 1.54f. Infected cells were counted and assessed for cytoophidia.

#### LCV formation

RAW 264.7 macrophages were seeded at 1 × 10^7^ cells per T75 flask one day prior to infection. At the day of the experiment, the cells were infected (MOI 50) with mCherry-producing *L. pneumophila* wild-type, Δ*icmT*, or Δ*ankX* (pNT102) grown to stationary phase (21 h in AYE), centrifuged (450 *g*, 10 min) to synchronize infection, and incubated at 37°C and 5% CO_2_ in a humidified incubator. The cells were washed once with DPBS (Gibco) 1 h post-infection and further incubated with fresh medium. At 2 h post-infection, cells were washed twice with DPBS (Gibco), then harvested by scraping. After centrifugation (450 *g*, 5 min, 4°C), cells were resuspended in homogenization buffer (20 mM HEPES pH 7.2, 250 mM sucrose, 0.5 mM EGTA) containing protease inhibitors (Pierce Protease Inhibitor Mini Tablets, EDTA-free; Thermo Fisher) and homogenized by passaging nine times through a ball homogenizer (Isobiotec) with an exclusion size of 8 µm. Samples (200 μL/well) were pipetted onto poly-L-lysine coated coverslips into 24-well plates and centrifuged (600 *g*, 10 min, 4°C), supernatants were removed, and samples were fixed with 4% PFA (30 min, RT). After fixation, coverslips were blocked with 1% BSA in DPBS for 1 h a RT, stained with rabbit anti-Atl3 antibody (Lubioscience: 16921-1-AP) diluted 1:25 in 1% BSA in DPBS. After 1 h incubation at RT, coverslips were washed in DPBS and incubated for another 1 h at RT with anti-rabbit Alexa 588 antibody (Life Technologies: A-11070) diluted 1:250 in 1% BSA in DPBS. Coverslips were washed again in DPBS, mounted onto microscope slides (Carl Roth) with ProLong Diamond Antifade Mountant with DAPI (Thermo Fisher), and dried overnight at RT.

Imaging was performed using a Leica SP8 DMi8 CS with AFC laser scanning microscope, an HC PL APO CS2 63×/1.4 oil objective, and Leica LAS X software (Leica). 30 images per tested condition were acquired at 10× zoom, and all bacteria were counted and assessed for Atl3-acquisition.

### LCV isolation and IMPDH2 binding assay

LCVs were purified from RAW 264.7 macrophages as previously described ^95^. One day prior to the experiment, the macrophages were seeded into four T75 flasks per *L. pneumophila* strain at a density of 1 × 10^7^ cells per flask to reach 80% confluency over-night. The cells were infected (MOI 50) with GFP-producing *L. pneumophila* wild-type or Δ*ankX* (pNT28) grown for 21 h in AYE to stationary phase. To synchronize infections, the cells were centrifuged (450 *g*, 10 min) and incubated for additional 50 min at 37°C and 5% CO_2_ in a humidified atmosphere. At 1 h post-infection, the cells were collected by scraping, washed three times with ice-cold DPBS (Gibco), centrifuged (450 *g*, 5 min, 4°C), and resuspended in homogenization buffer (20 mM HEPES pH 7.2, 250 mM sucrose, 0.5 mM EGTA) containing protease inhibitors (Pierce Protease Inhibitor Mini Tablets, EDTA-free; Thermo Fisher). Subsequently, infected cells were passaged nine times through a ball homogenizer (Isobiotec) with an exclusion size of 8 µm. Homogenates were blocked on ice with 2% FCS (30 min), incubated with an affinity-purified anti-SidC antibody (NeoMPS;^12^, 1:3,000, 1 h), and magnetic anti-rabbit IgG MicroBeads (130-048-602, Miltenyi Biotec, 1:25, 30 min). The LCVs were purified in a magnetic field using MACS-MS separation columns (Miltenyi Biotec). Homogenates were split equally onto four columns each, to test binding of purified IMPDH2 under different conditions, and columns were washed three times with 500 μL homogenization buffer. Purified IMPDH2 was diluted to 5 μM in phosphocholination buffer (20 mM Tris pH 7.0, 100 mM NaCl, 1 mM MgCl_2_, 1 mM DTT) and incubated at 25 °C for 6 h with ATP (1 mM), with ATP, CDP-choline (100 μM), and AnkX (0.1 μM), or none. Pre-incubated IMPDH2 was diluted to 1 μM in homogenization buffer and 100 μL were added to the isolated LCVs on the respective columns for 15 min at RT. As a control, one LCV sample per strain was incubated with 100 μL homogenization buffer. The flow-through was collected and analyzed by Western blot to control for equal passage of IMPDH2 through the column. Columns were washed again three times, then LCVs were eluted in 500 μL homogenization buffer, pelleted (500 *g*, 10 min), and resuspended in 100 μl SDS sample buffer for Western blot analysis.

LCVs were purified from *D. discoideum* Ax3 amoebae as previously described ^95^. Briefly, 1 × 10^7^ *D. discoideum* Ax3 producing calnexin-GFP (pAW016) were seeded into T75 flasks (3 flasks per condition) and grown overnight. The following day, the amoebae were infected (MOI 50) with DsRed-producing *L. pneumophila* JR32 or the Δ*sidM* mutant (pSW001) grown to stationary phase in AYE (21 h). Infection was synchronized by centrifugation (450 × *g*, 10 min, RT). After incubation at 25°C for 1 h, the cells were washed with SorC buffer (2 mM Na_2_HPO_4_, 15 mM KH_2_PO_4_, 50 µM CaCl_2_, pH 6.0), then harvested by scraping in homogenization (HS) buffer (20 mM HEPES, 250 mM sucrose, 0.5 mM EGTA, pH 7.2) containing a protease inhibitor cocktail tablet (Roche) ^96^. Subsequently, cells were homogenized using a ball homogenizer (Isobiotec) with an exclusion size of 8 µm. Homogenates were blocked with 2% FCS and the LCVs were isolated by immuno-magnetic separation and loaded onto four equilibrated MACS-MS columns as described above. The columns were washed three times with 0.5 ml HS buffer, incubated with 100 µL 1 µM AnkX, AnkX*, or LqsR, or with 100 µL HS buffer (15 min, RT), and washed again three times with 0.5 ml HS buffer. Finally, the LCVs were eluted in 0.5 ml HS buffer, centrifuged (500 × *g*, 10 min), and resuspended in 100 µL SDS sample buffer for Western blot analysis.

### Protein depletion by RNA interference and replication assay

For protein depletion experiments, 10 μM stock solutions of siRNA oligonucleotides (**Table S3**) or unspecific Allstars siRNA (Qiagen) were diluted 1:15 in RNAse-free water, and 3 μL were added per well of black 96-well plates (Corning). Each well was additionally supplemented with 24.25 μL RPMI without FCS mixed with 0.75 μL HiPerFect transfection reagent (Qiagen), and the plates were incubated at RT for 5-10 min. 2 × 10^4^ A549 cells per well were seeded in a volume of 175 μL on top of the transfection complexes and incubated for 48 h at 37°C and 5% CO_2_ in a humidified atmosphere. 48 h post-transfection, cells were infected (MOI 10) with GFP-producing *L. pneumophila* wild-type or Δ*icmT* (pNT28) as described above, and GFP-fluorescence was measured at 1 h and 24 h post-infection using a Cytation 5 Cell Imaging Multi-Mode Reader (BioTek).

To assess the efficiency of protein depletion, 9 μL of the 1:15 diluted siRNA (**Table S3**) was added to each well of a 24-well plate. Allstars siRNA (Qiagen) was used as a negative control.

72.75 μL RPMI 1640 without FCS mixed with 2.25 µL HiPerFect transfection reagent (Qiagen) was added to each well. After 5-10 min incubation at RT, 6×10^4^ A549 cells per well were added in a volume of 525 μL and transfected for 48 h at 37°C and 5% CO_2_ in a humidified incubator. Cells were harvested by trypsinization 48 h post-transfection, washed with ice-cold DPBS, and lysed in NP-40 buffer (20 mM Tris pH 8.0, 137 mM NaCl, 10% glycerol, 1% NP-40, 2 mM EDTA) supplemented with protease inhibitors (Pierce Protease Inhibitor Mini Tablets, EDTA-free; Thermo Fisher) for 30 min at 4°C on a rotating wheel (80 rpm). Soluble proteins were separated from cell debris by centrifugation (18,000 *g*, 4°C, 30 min) and analyzed by Western blot (see below).

### Scratch assay

To assess cell migration, A549 cells were seeded at a density of 4 × 10^4^ cells per well in 24-well plates, incubated for 48 h and infected (MOI 20) with GFP-producing *L. pneumophila* wild-type, Δ*icmT*, or Δ*ankX* (pNT28) grown in AYE to stationary phase. To synchronize infection, cells were centrifuged (450 *g*, 10 min) and incubated for 50 min at 37°C and 5% CO_2_ in a humidified incubator. Simultaneously to the infection, cells were treated with 10 µM MPA or 100 µM ribavirin, or left untreated. At 1 h post-infection, medium was replaced to remove extracellular bacteria, and a scratch was introduced to the cell layer in each well using the AutoScratch wound making tool (Agilent BioTek). The medium was replaced again to remove detached cells. Cells were imaged immediately after introducing the scratch, incubated at 37°C and 5% CO_2_ in a humidified incubator, and imaged again after 24 h. Imaging was performed using a Leica SP8 DMi8 CS with AFC laser scanning microscope with climate box at 37°C, an HC PL FLUOTAR 10×/0.3 air objective, white light laser, photomultiplier tube (PMT) detector, and Leica LAS X software (Leica). Tile-scans were acquired with 1× zoom to cover the area of the wound opening and merged with the Leica LAS X software. Wound areas were determined in ImageJ 1.54f by tracing the wound edges using a Wacom Intuos pen tablet (Wacom). For each condition, the wound closure was determined by comparing the wound opening after 24 h to the wound opening after 0 h.

To acquire representative images of the scratched cell monolayer, the untreated, infected and/or inhibitor-treated cells were fixed with 4% PFA for 20 min at 37°C, immediately after scratching the cell layers at 1 h post-infection, or 24 h after introducing the scratch wound. The fixed cells were washed once with DPBS and stored in DPBS at 4°C until imaging with a Nikon Eclipse Ti2, inverted widefield microscope with phase contrast. Images were acquired at 1× zoom with a Plan Fluor Ph1 DL 10×/0.3 air objective and a Nikon DS-Qi2 camera.

### Cytotoxicity assay of IMPDH2 inhibitor-treated cells

Cytotoxicity of IMPDH2 inhibitors MPA and ribavirin was assessed by live/dead staining and flow cytometric analysis of treated and untreated cells. To this end, 1·× 10^5^ A549 cells per well were seeded into 6-well plates. Two days after seeding, the medium of the cells was replaced by fresh medium (untreated control) or medium containing 10 µM MPA or 100 µM ribavirin and incubated further at 37°C and 5% CO_2_. After a total incubation time of 24 h, the supernatant of each well was collected separately, and the cells were harvested by trypsinization, combined with the respective supernatant, and stored on ice. A dead cell control was included by treating cells with sterile filtered 70% ethanol (in ddH_2_O) for 1 h prior to harvesting. The harvested cells were centrifuged (425 *g*, 5 min, 4°C) to remove the supernatant, washed twice with DPBS, and incubated with Zombie AquaTM dye (BioLegend, diluted 1:500 in DPBS) for 30 min in the dark. After the staining, cells were washed once with RPMI 1640 supplemented with FCS and once with DPBS, then fixed with 4% PFA for 1 h at RT. After fixation, cells were washed once with DPBS, resuspended in DPBS and stored at 4°C until analysis by flow cytometry. Dead cells (Zombie positive) were quantified using a Fortessa II flow cytometer and Diva software. The cell population was identified employing forward (FSC, 150 V) and sideward scatter (SSC, 150 V) gating, with a threshold of 200 each, and examined for the Zombie dye signal (Vio 525_50, 350 V). 10,000 events per sample were recorded. The data was analyzed with the software FlowJo. The Zombie positive cell population was gated using a dead control consisting of ethanol-treated cells as reference.

### Western blot analysis

For analysis of IMPDH2 binding to macrophage LCVs, protein samples were separated by SDS-PAGE (10% acrylamide gels), and transferred onto nitrocellulose membranes (Amersham) at 300 mA, 4°C for 90 min. Membranes were blocked (1 h, RT) with 3% milk in Tris-buffered saline containing 0.05% Tween 20 (TBST), and incubated with rabbit anti-IMPDH2 antibody (Abcam: ab131158), diluted 1:1,000 in fresh blocking solution, for 30 min at RT and overnight at 4°C. After overnight incubation, membranes were washed three times for 10 min with TBST and incubated for 1 h at RT with horseradish peroxidase (HRP)-conjugated secondary donkey anti-rabbit antibody (Cytiva: NA934-1ML) diluted 1:1,000 in TBST/3% milk. Membranes were washed another three times for 10 min with TBST, and peroxidase signal was detected with Westar Sun ECL Substrate 433 (Cyanagen) and the ImageQuant 800 (Amersham).

For the analysis of AnkX binding to *D. discoideum* LCVs, protein samples (15 µL) were run on 10% SDS-PAGE gels, transferred to nitrocellulose membranes, and blocked using Roti-Block (Carl Roth) 10-fold diluted in Tris-buffered saline containing 0.5% Tween20 (TBST) (for α-His and α-SidC Western blot) or 3% milk in TBST (α-LqsR Western blot). Subsequently, membranes were incubated with α-SidC (NeoMPS;^12^), α-His (Qiagen, 34670), or α-LqsR (Neosystem;^97^) antibodies at a dilution of 1:1,000 in the respective blocking solution for 30 min at RT and overnight at 4°C. Primary antibodies were detected using HRP-coupled sheep α-mouse or donkey α-rabbit antibodies (GE Healthcare) and Westar Sun (Cyanagen) chemiluminescent substrate.

To assess phosphocholination by Western blot, protein samples were separated by SDS-PAGE (10% acrylamide gels) and transferred onto polyvinylidene fluoride (PVDF) membrane (Amersham) at 300 mA, 4°C for 90 min. PVDF membranes were blocked with 10% Roti-Block (Carl-Roth) in TBST for 1 h at RT, and probed with an anti-phosphocholine antibody (1:1,000; Sigma-Aldrich: M1421-1MG, clone TEPC 15) for 30 min at RT and overnight at 4°C. Membranes were further processed as described above using a sheep anti-mouse HRP-conjugated secondary antibody (1:1,000, Cytiva: NA931-1ML).

To assess depletion of IMPDH2 in cells treated with siRNA oligonucleotides, cell lysate samples were separated by SDS-PAGE (10% acrylamide gels), and blotted onto nitrocellulose membranes at 300 mA, 4°C for 90 min. Membranes were blocked with 3% milk in TBST for 1 h at RT, and incubated with rabbit anti-IMPDH2 (1:1,000, Abcam: ab131158), rabbit anti-Hsp90 (1:500, Abcam: ab19021), or rabbit anti-GAPDH (1:1,000, Cell Signaling Technology: 2118) for 30 min at RT and overnight at 4°C. Washing steps, incubation with secondary antibody and imaging were performed as described above.

### Western Blot for IMPDH2 PCylation kinetics

Recombinant IMPDH2 or Rab1b (5 µM) was incubated with CDP-choline (100 µM) and either no nucleotide, ATP, or GTP (100 µM) in 20 mM HEPES, 100 mM NaCl, 1 mM MgCl_2_, 1 mM DTT, pH 7 for 15 min at RT to allow nucleotide binding. Reactions were initiated by adding His-GFP-AnkX (500 nM) and sampled at 0, 30, 60, 120, (180), 240, 360, (480), and 1440 min. The phosphocholination reaction was stopped at the time points indicated by adding SDS-PAGE loading buffer. Proteins (500 ng per lane) were separated by 12% SDS-PAGE and transferred to MeOH-activated PVDF membranes (Immobilon®, Merck Millipore) using a semi-dry blotter (Blotting buffer: 48 mM Tris, 39 mM glycine, 1.3 mM SDS, 20% methanol). Membranes were washed in TBST for 5 min, blocked with 1× Roti®-Block (Carl Roth) in TBST (0.1% Tween20) for 1 h at RT, and incubated overnight at 4°C with primary antibody (TEPC-15, Sigma, 1:1,000). After three 15 min washes in TBST, membranes were incubated with HRP-conjugated secondary antibody (goat-anti-mouse IgG, Thermo Fisher) for 1 h, washed again, and developed using SuperSignal™ West Dura (Thermo Fisher). Chemiluminescence was detected with an Intas ECL Chemocam (Intas Science Imaging Instruments). Experiments were performed in three biological replicates. Band intensities were quantified using ImageJ 1.45g, with the 24 h sample set to 100% and all other bands expressed relative to this value.

To verify equal protein loading across samples and ensure that subsequent antibody-based detection was not influenced by differences in protein amounts, membranes were subjected to colloidal silver staining. Membranes were washed three times with deionized water prior to staining. A colloidal silver staining solution was freshly prepared for each experiment (15–25 mL per membrane). The solution consisted of 0.4 g ferrous sulfate heptahydrate dissolved in 47 mL deionized water, followed by the addition of 2.5 mL of 40% sodium citrate dihydrate. Subsequently, 0.5 mL of 20% silver nitrate stock solution was added dropwise until a dark brown precipitate formed. The mixture was vortexed for approximately 1 min. Membranes were incubated in the staining solution for 10 min under gentle agitation. Staining was terminated by thoroughly washing the membranes in deionized water.

## Statistical analysis

If not indicated otherwise, all statistical analysis were performed with GraphPad prism version 9.5.1, GraphPad Software Inc (www.graphpad.com). One-way ANOVA with post hoc Tukey test and two-way ANOVA with post hoc Dunnett’s test were performed to evaluate significant differences between experimental conditions and controls. Significance levels are indicated as *, **, ***, or **** and represent probability values of less than 0.5, 0.01, 0.001, or 0.0001, respectively. The value “n” represents the number of cells analyzed per condition.

## Data availability

The mass spectrometry proteomics data have been deposited to the ProteomeXchange Consortium via PRIDE ^98^ partner repository with the dataset identifier PXD059102 or via MassIVE (MSV000099455) and identifier PXD069380. All other data generated in this study are included in the manuscript or in the supplemental information.

### Use of artificial intelligence

Text editing and language polishing were performed using the artificial intelligence (AI) language model ChatGPT (OpenAI), version GPT-5. The authors reviewed and revised all text generated by the model and take full responsibility for the content and accuracy of the manuscript. No data, results, analyses or figures were generated or interpreted by the AI.

## Acknowledgements

P.S. and H.H. thank Jacqueline Cherfils and Pavlina Dubois for help with the purification of AnkX. We acknowledge the Functional Genomics Center Zurich (FGCZ) of the University of Zurich and ETH Zurich, and in particular Serge Chesnov, Sibylle Pfammatter, and Peter Gehrig, for their scientific and technical support on proteomics analysis. Transmission electron microscopy and confocal laser scanning microscopy were performed with the equipment provided and maintained by the Center for Microscopy and Image Analysis (ZMB), University of Zurich. Research performed in the laboratory of H.H. was supported by the Swiss National Science Foundation (SNSF; project grant 207826) and the Vontobel Foundation (stipend awarded to P.S.). M.S.K. and A.I. acknowledge funding by the Deutsche Forschungsgemeinschaft (DFG; RTG2771). M.S.K. and A.I. acknowledge technical support from the SPC facility at EMBL Hamburg and access to the core facilities and laboratories of the Centre for Structural Systems Biology (CSSB, Hamburg).

## Author Contributions

Conceptualization: PS, MSK, ALS, CH, AI, HH

Methodology: PS, MSK, ALS, PO, BS, VP, AB, CS, PJ

Investigation: PS, MSK, ALS, PO, BS, VP, DFO, AB, CS

Visualization: PS, MSK, ALS, PO

Funding acquisition: PS, HS, CH, AI, HH

Project administration: HH

Supervision: PJ, HS, CH, AI, HH

Writing – original draft: PS, MSK, HH

Writing – review/editing: PS, MSK, ALS, PO, BS, VP, DFO, AB, CS, PJ, HS CH, AI, HH

## Competing interests

The authors declare no conflict of interest.

## Abbreviations

DLS: dynamic light scattering
EM: electron microscopy
Icm/Dot: intracellular multiplication/ defective organelle trafficking
IMPDH2: inosine monophosphate dehydrogenase type II
LCV: *Legionella*-containing vacuole
MPA: mycophenolic acid
PC: phosphocholine
T4SS: type IV secretion system.

---

**Figure S1.**
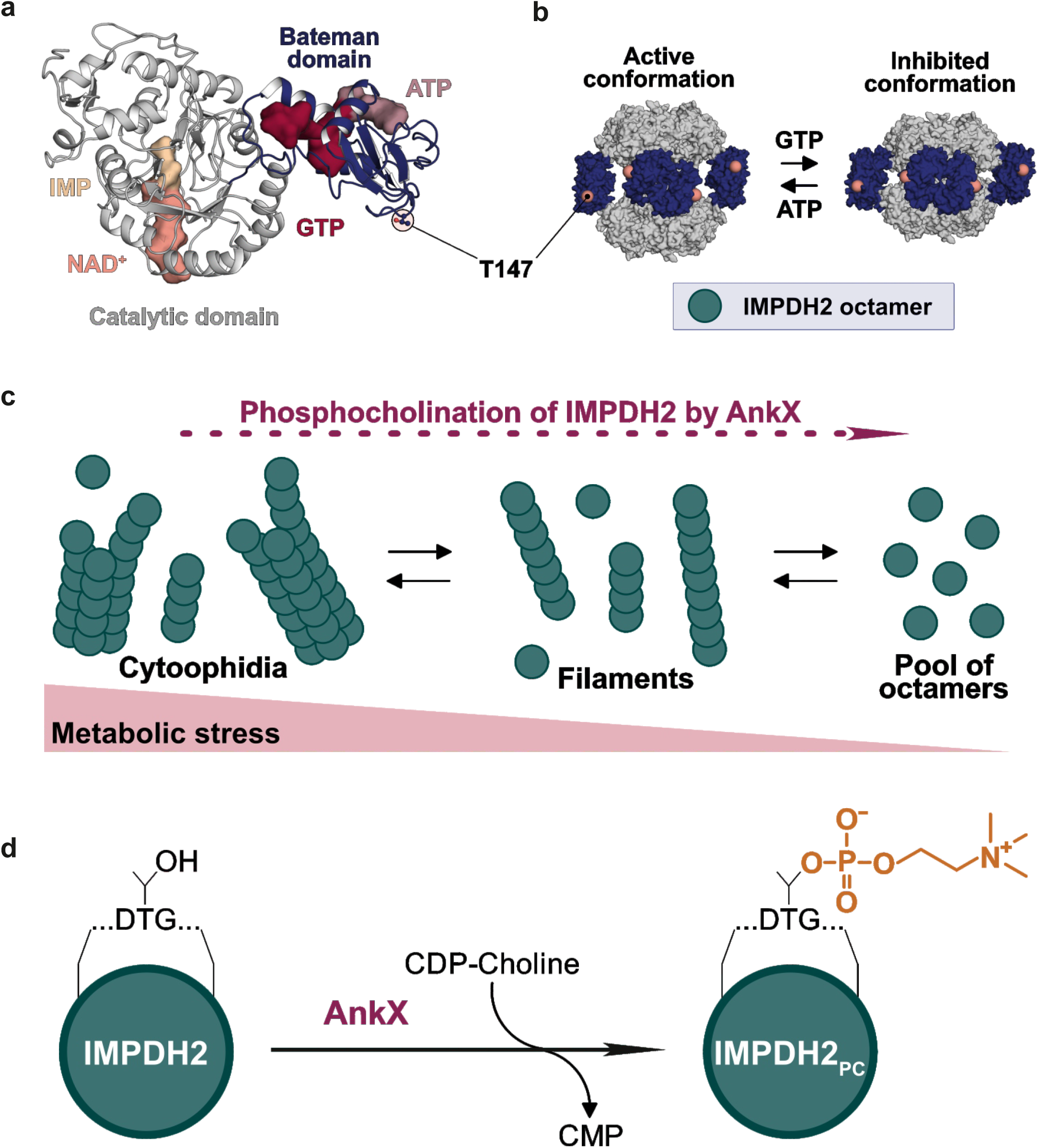
Structure, oligomerization, and activity of IMPDH2. (**a**) Cartoon depiction of the IMPDH2 monomer in its GTP-bound, inhibited state (PDB ID: 6U9O). The catalytic domain (grey) is bound to IMP (gold) and NAD^+^ (salmon), the regulatory domain (Bateman domain, blue) comprises three nucleotide binding pockets for ATP (lavender) and GTP (red). Bound nucleotides are depicted as surface-representation and the phosphocholination site T147_IMPDH2_, located in the regulatory domain, is depicted as ball-and-stick-representation and colored according to atom identity. (**b**) In solution, IMPDH2 can form an octamer upon binding of ATP or GTP. The shape of the octamer is determined by the combination of bound nucleotides, with ATP binding inducing the active, bowed conformation (PDB ID: 6U8N), and GTP binding inducing the compressed, inhibited conformation (PDB ID: 6U9O). The location of T147_IMPDH2_ in the individual monomers of the octamers is depicted as a salmon-colored sphere. Both octamer forms can assemble into filaments and higher molecular structures called cytoophidia. (**c**) Model of IMPDH2 assembly dynamics under metabolic stress, in which equilibrium between free octamers, filaments and cytoophidia is regulated by cellular GTP demand. (**d**) The phosphocholination of IMPDH2 by the *L. pneumophila* effector AnkX shifts the equilibrium toward free octamers, preventing filament and cytoophidia formation. Filament disassembly was detected *in vitro* and cytoophidia disappearance was observed *in cellulo* (based on ^42^, and https://www.biorxiv.org/content/10.1101/2024.07.29.605679v2).

**Figure S2.**
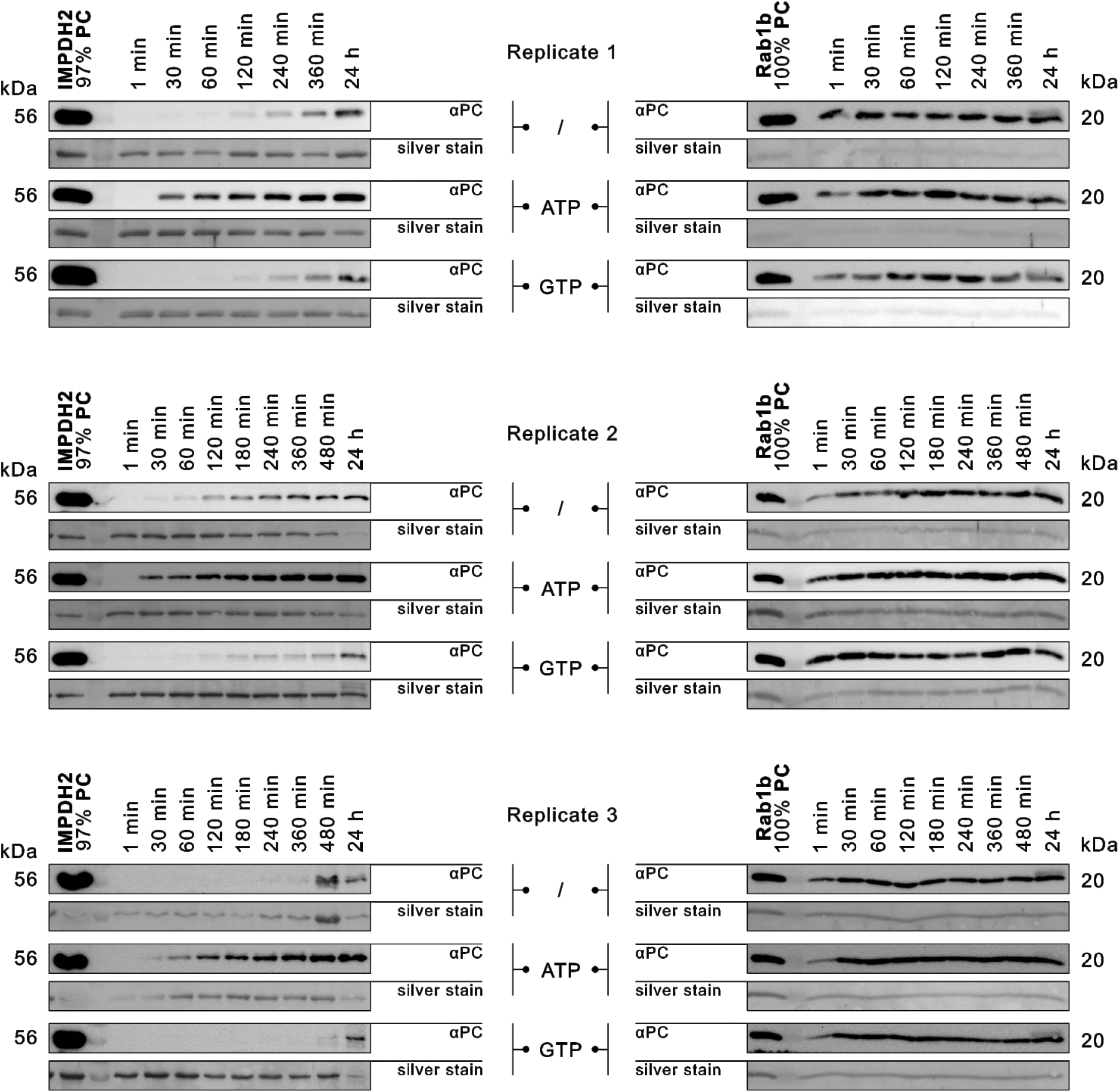
Full set of Western blots. Independent triplicates of Western blots are shown (Fig. 1d).

**Figure S3.**
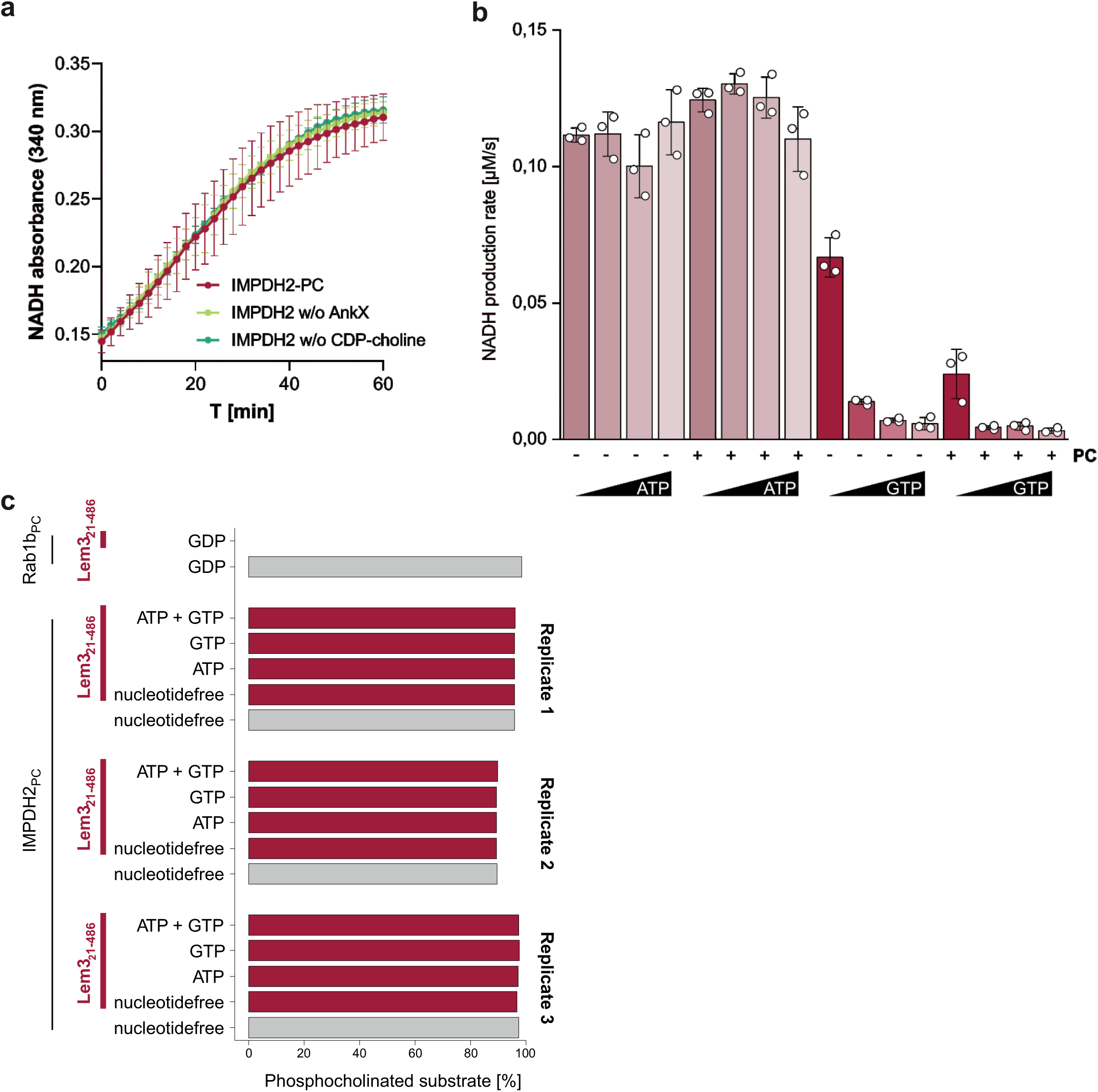
Catalytic activity of IMPDH2 and Lem3. (**a**) IMPDH2 was incubated for 3 h with CDP-choline and AnkX for phosphocholination, or – as controls – only with CDP-choline or AnkX. IMPDH2 activity (NAD⁺-dependent conversion of IMP to XMP) was quantified measuring NADH production at the time points indicated. (**b**) IMPDH2 activity for phosphocholinated and unmodified IMPDH2 in absence and presence of different concentrations of ATP or GTP (0.2, 1, 5, 25 mM) was quantified measuring NADH production over time. Means (±SD) represent three independent biological replicates (unpaired, two-tailed t-test). (**c**) Phosphocholinated Rab1b (Rab1b_PC_) and IMPDH2 (IMPDH2_PC_) were incubated (nucleotide loading status as indicated) with Lem3_21-486_ (24 h, RT) and analyzed by iMS. Dephosphocholination can only be detected for Rab1b_PC_. Three independent biological replicates are shown for IMPDH2.

**Figure S4.**
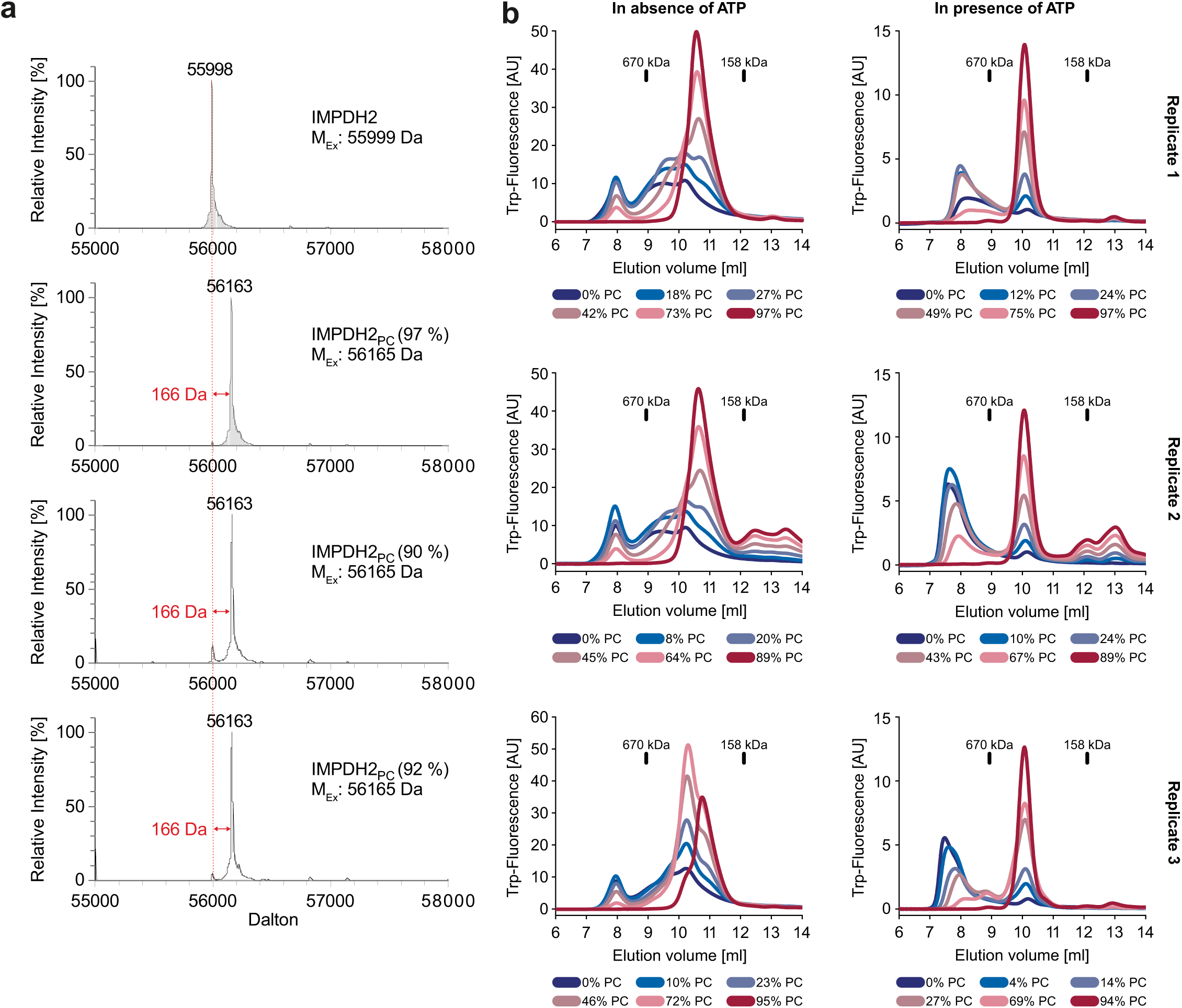
Analysis of IMPDH2 modification by AnkX and effects on IMPDH2 filament formation. (**a**) iMS of IMPDH2 and biological triplicates of preparative IMPDH2_PC_.The spectra were taken from the same samples used for aSEC (Fig. 2e), DLS (Fig. 2f), and LC-MS/MS for identification of the modification site (Fig. 3a-c). (**b**) aSEC of unmodified and phosphocholinated IMPDH2 in absence or presence of ATP. Purified IMPDH2 (50 μg) phosphocholinated to varying percentages (indicated below the respective graph) was loaded and detected using tryptophan fluorescence (excitation at 280 nm, emission at 340 nm). Elution volumes of reference proteins are indicated (thyreoglobulin: 670 kDa, γ-globulin:158 kDa). Three independent biological replicates are shown.

**Figure S5.**
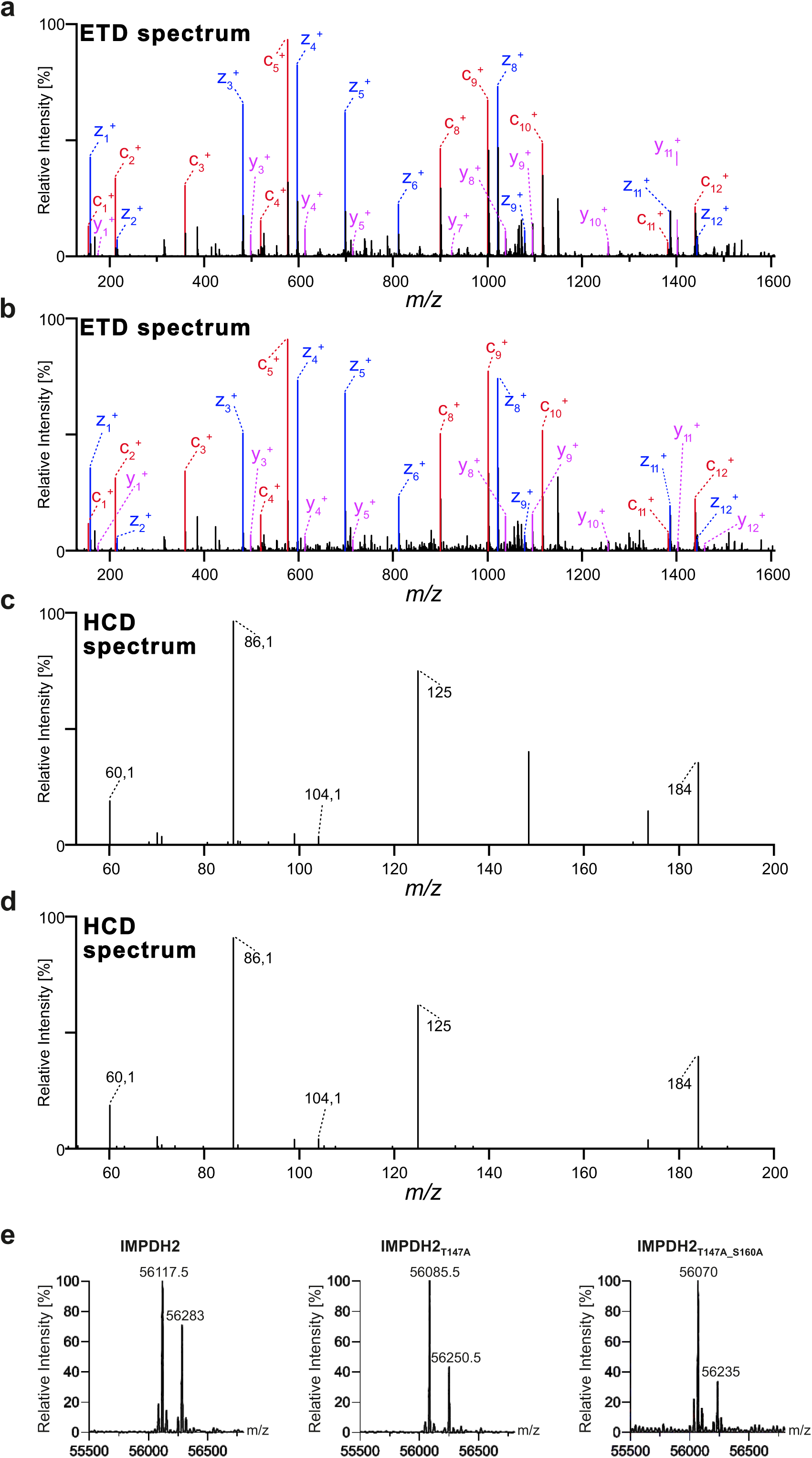
Fragmentation spectra of phosphocholinated IMPDH2 peptide. The IMPDH2 peptide HGFCGIPITDTGR (m/z 532.5854) analyzed by (**a**, **b**) ETD and (**c**, **d**) HCD MS. ETD fragmentation spectra (MS2) with annotated c, y, and z ions of biological replicates of (**a**) IMPDH2_PC_ (89%) and (**b**) IMPDH2_PC_ (95%). Ions identified are marked in red, blue and magenta. (**c**, **d**) HCD fragmentation spectra (MS3) of the cleaved-off phosphocholine from biological replicates of (**c**) IMPDH2_PC_ (89%) and (**d**) IMPDH2_PC_ (95%). (**e**) iMS of 5 µM IMPDH2, IMPDH2_T147A_, or IMPDH2_T147A_S160A_ after 1 h incubation with 1 mM ATP, 100 µM CDP-choline, and 0.5 µM AnkX at 25°C.

**Figure S6.**
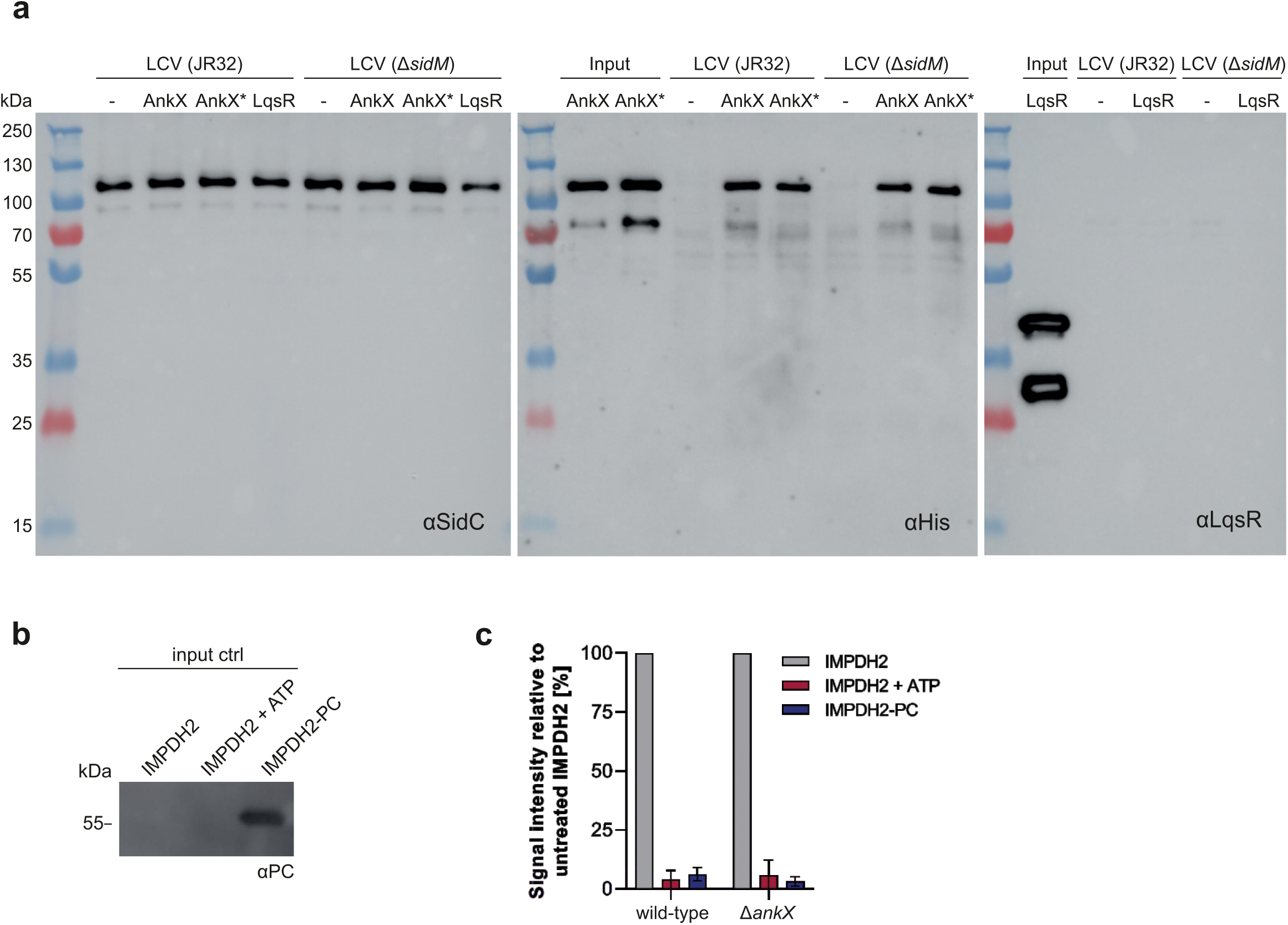
AnkX, AnkX_H229A_, and IMPDH2 bind to purified LCVs. (**a**) LCVs were isolated from homogenized *D. discoideum* infected with *L. pneumophila* wild-type or Δ*sidM* (lacking the Rab1 guanine nucleotide exchange factor SidM). The LCVs bound to separation columns were washed and incubated with 1 µM wild-type AnkX, catalytically inactive AnkX* (AnkX_H229A_), the response regulator LqsR, or with 100 µl buffer. After washing, the LCV samples were eluted and analyzed by Western blot with the following antibodies: α-SidC (SidC, 111 kDa; LCV preparation control), α-His (His-AnkX or His-AnkX*; 110.5 kDa), or α-LqsR (LqsR, 39.4 kDa and ∼30 kDa fragment ^97^; negative control). As input controls, 4.5 pmol purified His-AnkX, His-AnkX*, and 3C-protease treated His-LqsR was used. (**b**) Phosphocholination of untreated IMPDH2 (IMPDH2), or incubated with ATP (IMPDH2 + ATP), or ATP, AnkX_1-800_, and CDP-choline (IMPDH2_PC_) was assessed by Western blot analysis using an anti-phosphocholine antibody (αPC) and used for LCV binding experiments. (**c**) Western blots of 3 independent experiments assessing IMPDH2 binding to the LCV were analyzed by densitometry. Means and STD of four biological replicates are shown.

**Figure S7.**
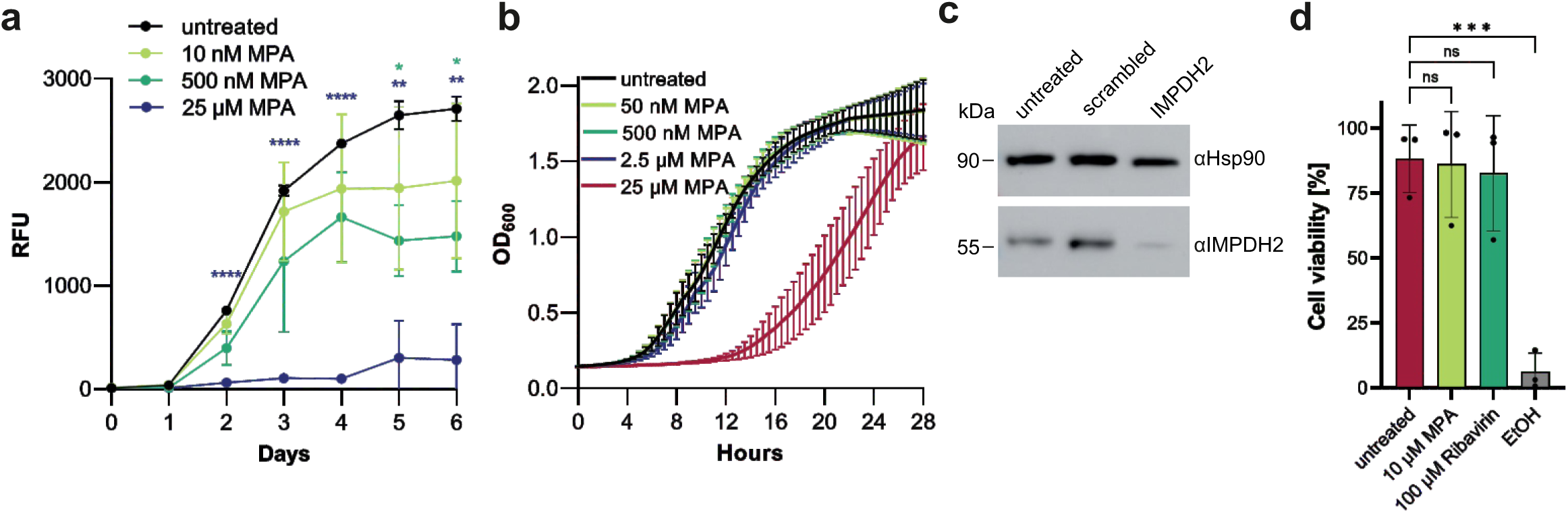
IMPDH2 promotes growth of *L. pneumophila* in macrophages. (**a**) RAW 264.7 macrophages were treated with MPA at the indicated concentrations and infected (MOI 1) with wild-type *L. pneumophila* producing GFP (pNT28), and intracellular replication at 37°C was assessed daily by relative fluorescent units (RFU). For each time point, the difference of the MPA-treated cells to the untreated cells was tested by a two-way ANOVA with Dunnett’s multiple comparisons test; *, P < 0.05; **, P < 0.01; ***, P < 0.001; ****, P < 0.0001. Means and SD of technical triplicates are shown. (**b**) To assess the toxicity of MPA for *L. pneumophila*, the growth of strain JR32 in AYE medium starting at an OD of 0.1 in the absence or presence of the indicated MPA concentrations was assessed by OD_600_ in intervals of 30 min. Means and SD of three independent experiments are shown. (**c**) To assess the depletion efficiency of siRNA oligonucleotides targeting IMPDH2, A549 cells were treated for 48 h with a combination of four different oligonucleotides, with AllStars unspecific oligonucleotides (“scrambled”) as a control for off-target effects or left untreated. Western blot analysis was performed using the antibodies indicated (αHsp90, αIMPDH2). Hsp90 was used as a loading control. Data is representative of three independent experiments. (**d**) To assess the toxicity of MPA or ribavirin for A549 cells, the cells were treated for 24 h with 10 µM MPA or 100 µM ribavirin, stained with Zombie Aqua™ dye, and fixed with 4% PFA. Cytotoxicity was assessed by quantifying the uptake of the dye by flow cytometry (excitation 405 nm). Viability of inhibitor-or ethanol-treated cells (dead control) was compared to untreated cells by one-way ANOVA with Dunnett’s multiple comparisons test; ***, P < 0.001. Shown are means and SD of three independent experiments.

**Figure S8.**
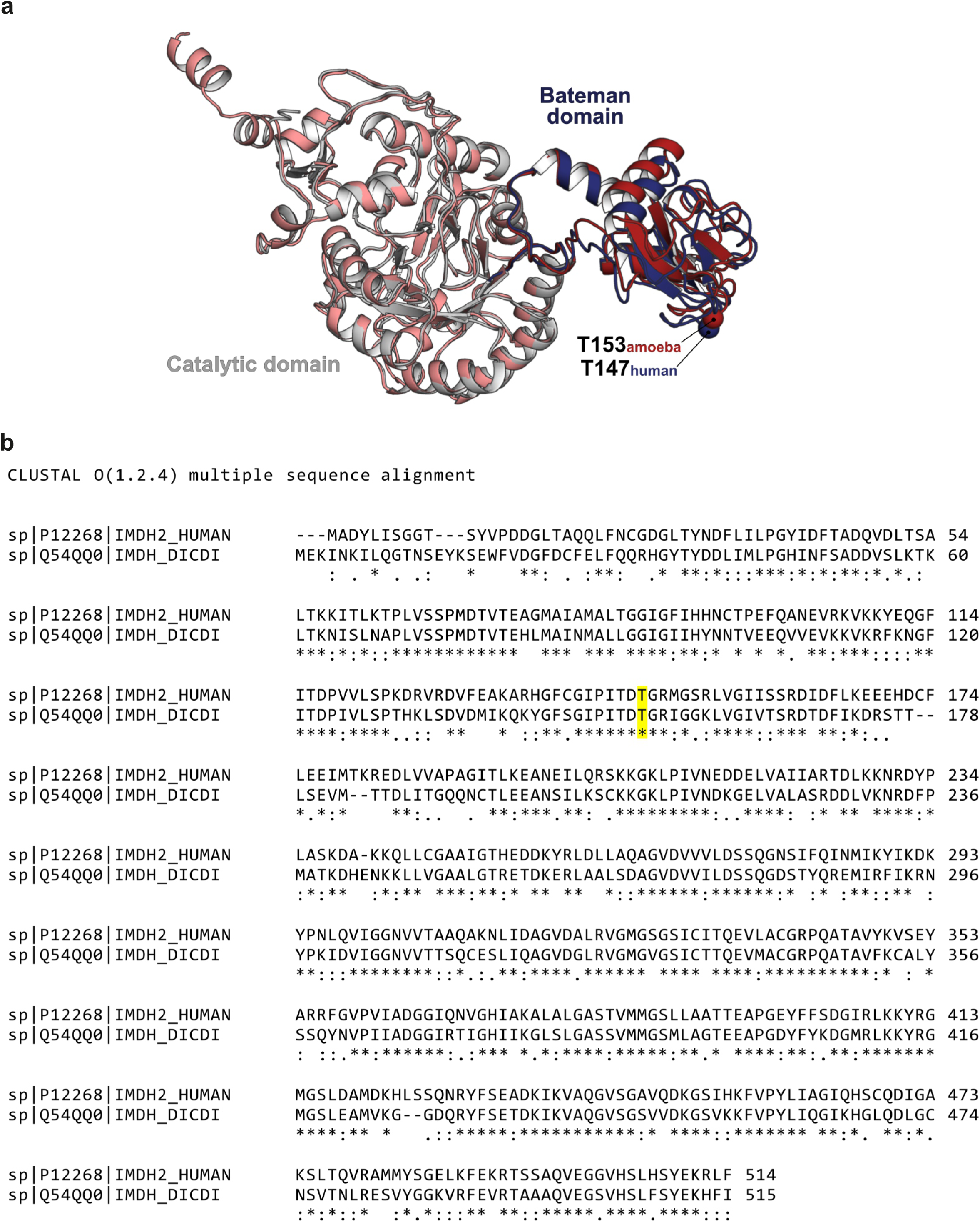
Structural and sequence comparison of human and amoeba IMPDH. **(a**) Superposition of the cryo-EM structure of human IMPDH2 (PDB ID: 6U9O) and the AlphaFold-predicted structure of *D. discoideum* IMPDH (AF-Q54QQ0-F1-model_v6). In the human enzyme, the catalytic domain is shown in grey and the Bateman domain in blue; in the amoeba model, the catalytic domain is shown in nude and the Bateman domain in red. Domain labels correspond to the colour scheme used for the human IMPDH2 structure. The Cα atoms of the conserved threonine residues (T147 in human IMPDH2 and T153 in *D. discoideum* IMPDH) are depicted as spheres. (**b**) Multiple sequence alignment of human (UniProt P12268) and *D. discoideum* (UniProt Q54QQ0) IMPDH generated using Clustal Omega (v1.2.4). The conserved threonine residues (T147 in human and T153 in amoeba) are highlighted.

**Table S1.**
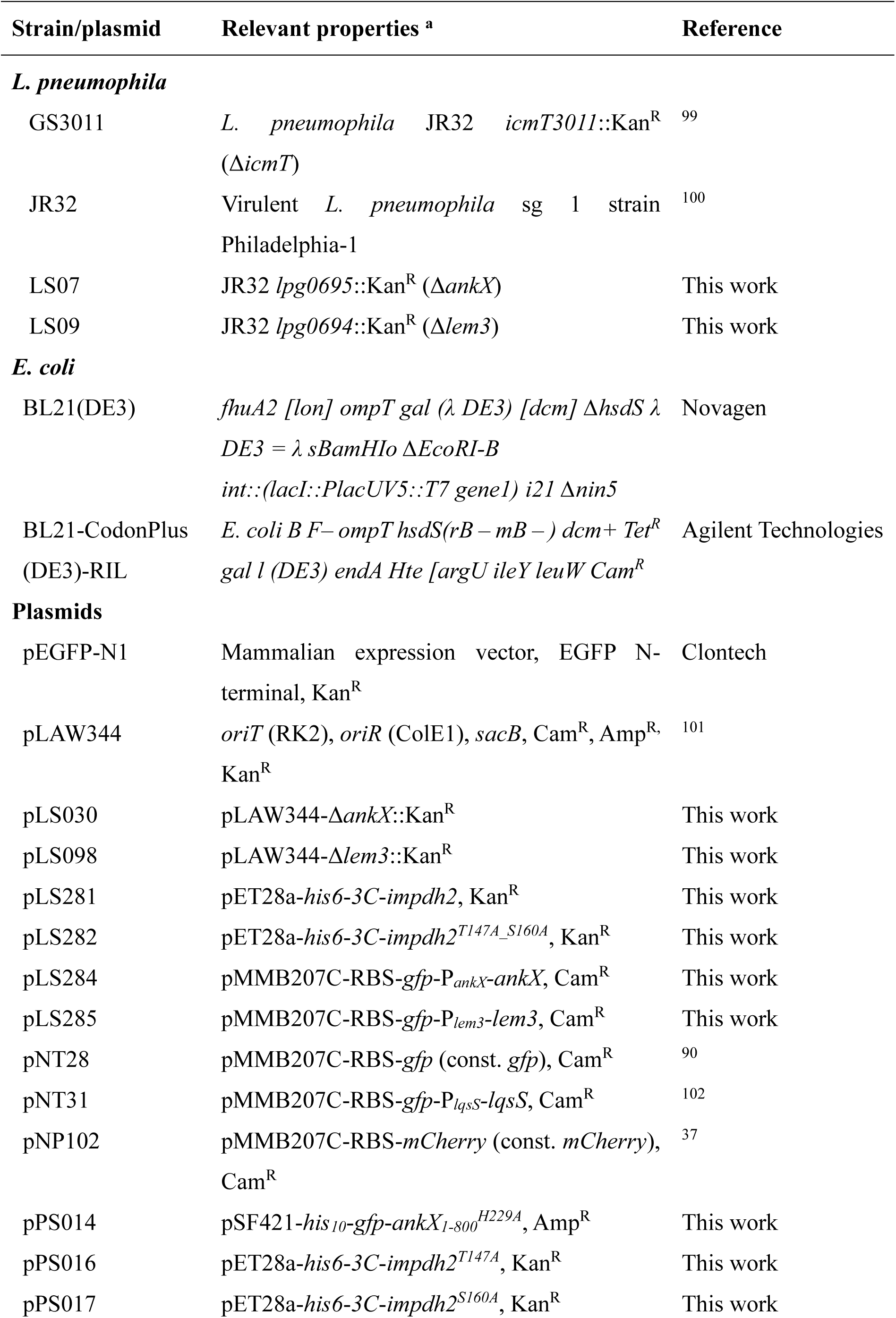
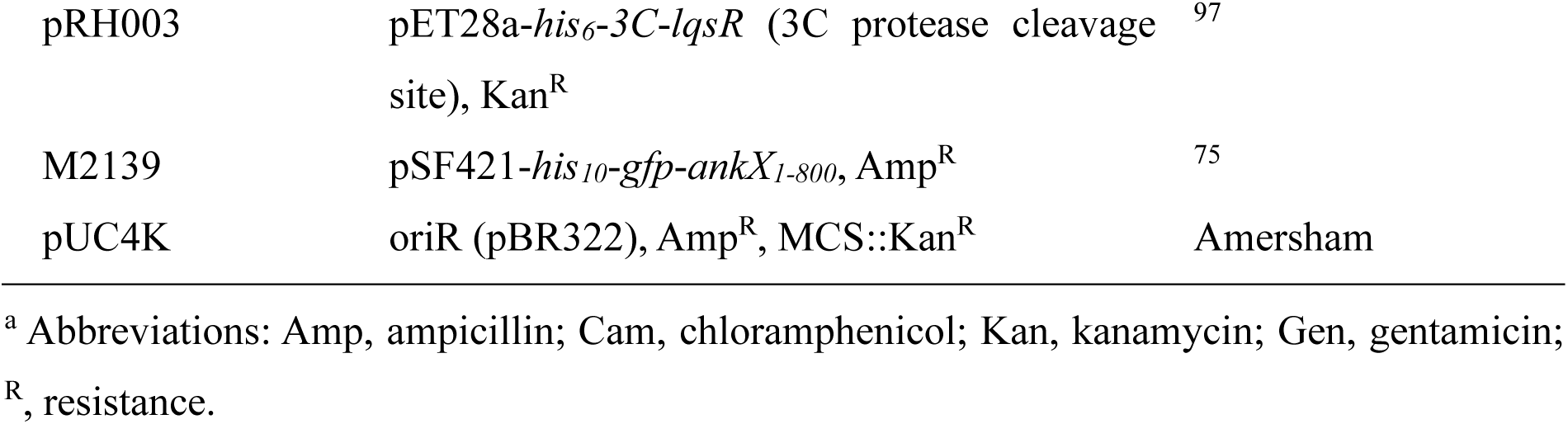
Bacterial strains and plasmids used in this study.

**Table S2.**
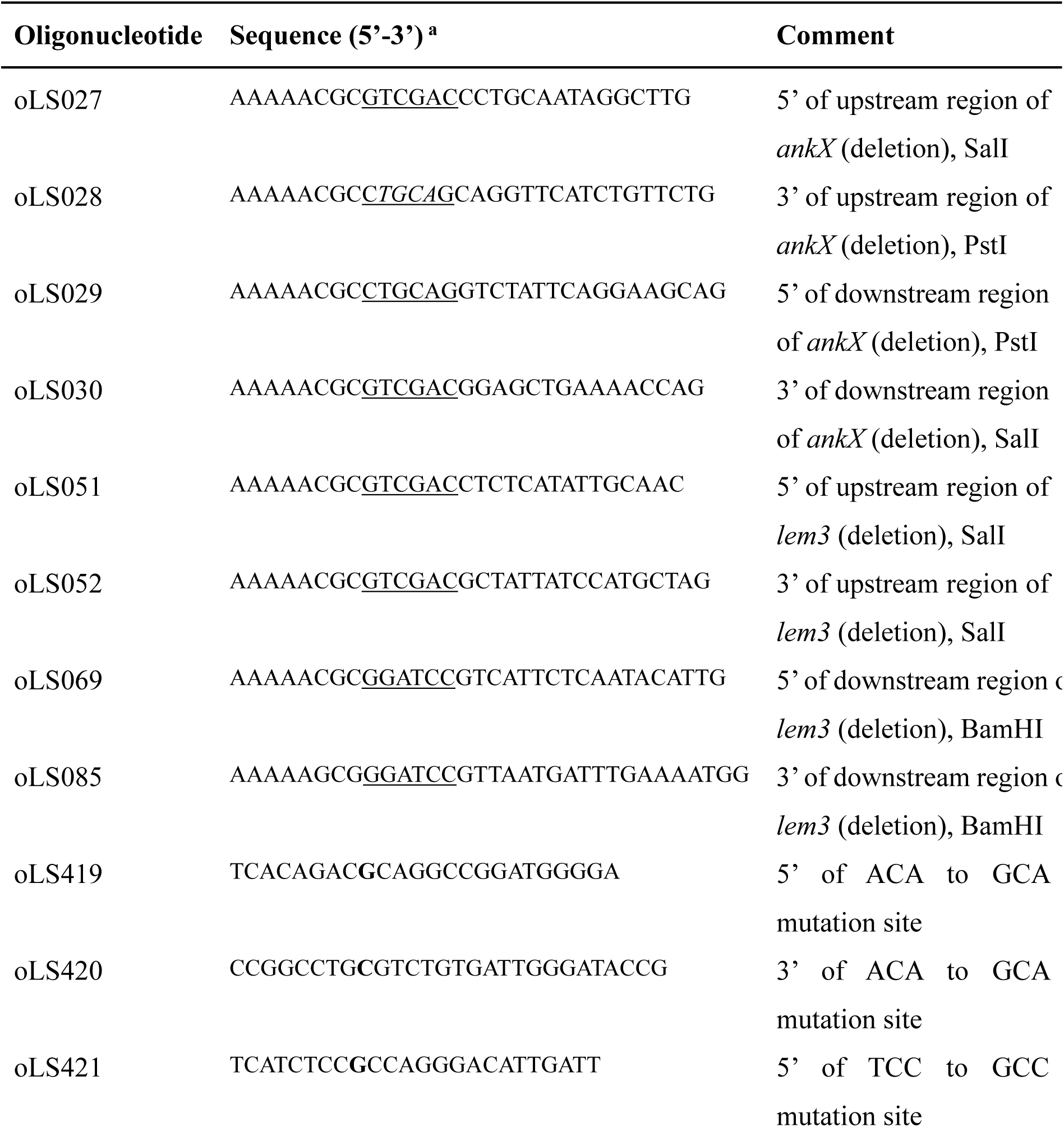
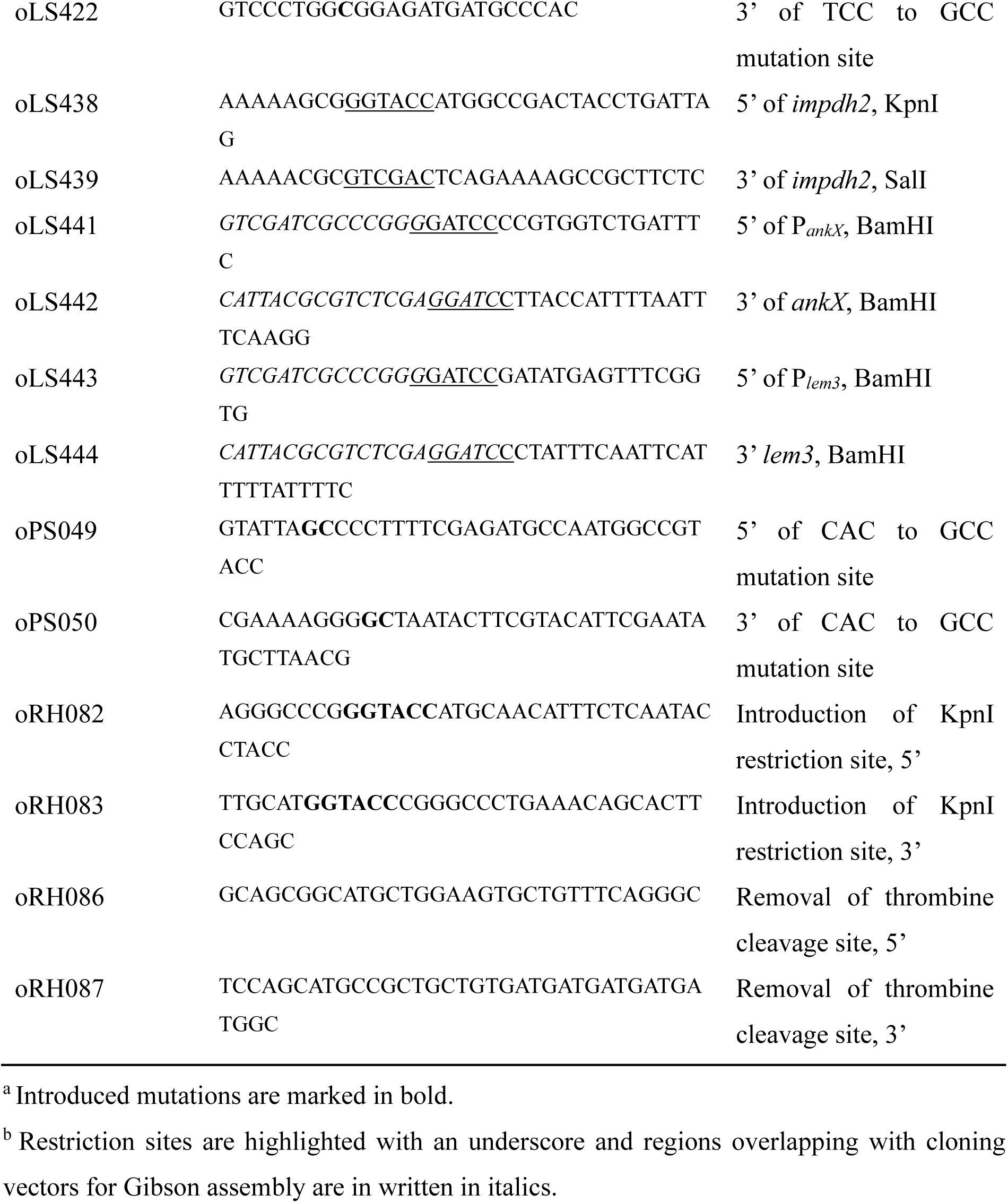
Oligonucleotides used in this study.

**Table S3.** Oligonucleotides used for RNA interference.

| NCBI gene | Gene description | Entrez Gene ID | Product name | Product ID |
| --- | --- | --- | --- | --- |
|  | AllStars | 0 | Unspecific_AllStars<br>_1 | SI03650318 |
| ARF1 | ADP-ribosylation<br>factor 1 | 375 | Hs_ARF1_10 | SI02757272 |
| IMPDH2 | Inosine<br>monophosphate<br>dehydrogenase 2 | 3615 | Hs_IMPDH2_5 | SI00300552 |
| IMPDH2 | Inosine<br>monophosphate<br>dehydrogenase 2 | 3615 | Hs_IMPDH2_6 | SI02780610 |
| IMPDH2 | Inosine<br>monophosphate<br>dehydrogenase 2 | 3615 | Hs_IMPDH2_7 | SI03031189 |
| IMPDH2 | Inosine<br>monophosphate<br>dehydrogenase 2 | 3615 | Hs_IMPDH2_8 | SI03037440 |

## References

1 Newton, H. J., Ang, D. K., van Driel, I. R. & Hartland, E. L. Molecular pathogenesis of infections caused by *Legionella pneumophila*. Clin Microbiol Rev 23, 274–298 (2010).

2 Mondino, S. et al. Legionnaires’ disease: state of the art knowledge of pathogenesis mechanisms of *Legionella*. Ann Rev Pathol 15, 439–466 (2020).

3 Hilbi, H. & Buchrieser, C. Microbe profile: *Legionella pneumophila* - a copycat eukaryote. Microbiology (Reading) 168, doi: 10.1099/mic.1090.001142 (2022).

4 Romanov, K. A. & O’Connor, T. J. *Legionella pneumophila*, a Rosetta stone to understanding bacterial pathogenesis. J Bacteriol 206, e0032424 (2024).

5 Kubori, T. & Nagai, H. The type IVB secretion system: an enigmatic chimera. Curr Opin Microbiol 29, 22–29 (2016).

6 Qiu, J. & Luo, Z. Q. *Legionella* and *Coxiella* effectors: strength in diversity and activity. Nat Rev Microbiol 15, 591–605 (2017).

7 Lockwood, D. C., Amin, H., Costa, T. R. D. & Schroeder, G. N. The *Legionella pneumophila* Dot/Icm type IV secretion system and its effectors. Microbiology (Reading) 168, doi: 10.1099/mic.1090.001187 (2022).

8 Finsel, I. et al. The *Legionella* effector RidL inhibits retrograde trafficking to promote intracellular replication. Cell Host Microbe 14, 38–50 (2013).

9 Bärlocher, K. et al. Structural insights into *Legionella* RidL-Vps29 retromer subunit interaction reveal displacement of the regulator TBC1D5. Nat Commun 8, 1543 (2017).

10 Romano-Moreno, M. et al. Molecular mechanism for the subversion of the retromer coat by the *Legionella* effector RidL. Proc Natl Acad Sci U S A 114, E11151–E11160 (2017).

11 Yao, J. et al. Mechanism of inhibition of retromer transport by the bacterial effector RidL. Proc Natl Acad Sci U S A 115, E1446–E1454 (2018).

12 Weber, S. S., Ragaz, C., Reus, K., Nyfeler, Y. & Hilbi, H. *Legionella pneumophila* exploits PI(4)*P* to anchor secreted effector proteins to the replicative vacuole. PLoS Pathog 2, e46 (2006).

13 Luo, X. et al. Structure of the *Legionella* virulence factor, SidC reveals a unique PI(4)*P*-specific binding domain essential for its targeting to the bacterial phagosome. PLoS Pathog 11, e1004965 (2015).

14 Brombacher, E. et al. Rab1 guanine nucleotide exchange factor SidM is a major phosphatidylinositol 4-phosphate-binding effector protein of *Legionella pneumophila*. J Biol Chem 284, 4846–4856 (2009).

15 Schoebel, S., Blankenfeldt, W., Goody, R. S. & Itzen, A. High-affinity binding of phosphatidylinositol 4-phosphate by *Legionella pneumophila* DrrA. EMBO Rep 11, 598–604 (2010).

16 Zhu, Y. et al. Structural mechanism of host Rab1 activation by the bifunctional *Legionella* type IV effector SidM/DrrA. Proc Natl Acad Sci U S A 107, 4699–4704 (2010).

17 Del Campo, C. M. et al. Structural basis for PI(4)P-specific membrane recruitment of the *Legionella pneumophila* effector DrrA/SidM. Structure 22, 397–408 (2014).

18 Hammond, G. R., Machner, M. P. & Balla, T. A novel probe for phosphatidylinositol 4-phosphate reveals multiple pools beyond the Golgi. J Cell Biol 205, 113–126 (2014).

19 Müller, M. P. et al. The *Legionella* effector protein DrrA AMPylates the membrane traffic regulator Rab1b. Science 329, 946–949 (2010).

20 Bhogaraju, S. et al. Phosphoribosylation of ubiquitin promotes serine ubiquitination and impairs conventional ubiquitination. Cell 167, 1636–1649 (2016).

21 Qiu, J. et al. Ubiquitination independent of E1 and E2 enzymes by bacterial effectors. Nature 533, 120–124 (2016).

22 Kotewicz, K. M. et al. A single *Legionella* effector catalyzes a multistep ubiquitination pathway to rearrange tubular endoplasmic reticulum for replication. Cell Host Microbe 21, 169–181 (2017).

23 Mukherjee, S. et al. Modulation of Rab GTPase function by a protein phosphocholine transferase. Nature 477, 103–106 (2011).

24 Tan, Y., Arnold, R. J. & Luo, Z. Q. *Legionella pneumophila* regulates the small GTPase Rab1 activity by reversible phosphorylcholination. Proc Natl Acad Sci U S A 108, 21212–21217 (2011).

25 Goody, P. R. et al. Reversible phosphocholination of Rab proteins by *Legionella pneumophila* effector proteins. EMBO J 31, 1774–1784 (2012).

26 Kaspers, M. S. et al. Dephosphocholination by *Legionella* effector Lem3 functions through remodelling of the switch II region of Rab1b. Nat Commun 14: 2245 (2023).

27 Isberg, R. R., O’Connor, T. J. & Heidtman, M. The *Legionella pneumophila* replication vacuole: making a cosy niche inside host cells. Nat Rev Microbiol 7, 13–24 (2009).

28 Hubber, A. & Roy, C. R. Modulation of host cell function by *Legionella pneumophila* type IV effectors. Ann Rev Cell Dev Biol 26, 261–283 (2010).

29 Steiner, B., Weber, S. & Hilbi, H. Formation of the *Legionella*-containing vacuole: phosphoinositide conversion, GTPase modulation and ER dynamics. Int J Med Microbiol 308, 49–57 (2018).

30 Garcia-Rodriguez, F. J., Buchrieser, C. & Escoll, P. *Legionella* and mitochondria, an intriguing relationship. Int Rev Cell Mol Biol 374, 37–81 (2023).

31 Urwyler, S. et al. Proteome analysis of *Legionella* vacuoles purified by magnetic immunoseparation reveals secretory and endosomal GTPases. Traffic 10, 76–87 (2009).

32 Hoffmann, C. et al. Functional analysis of novel Rab GTPases identified in the proteome of purified *Legionella*-containing vacuoles from macrophages. Cell Microbiol 16, 1034–1052 (2014).

33 Naujoks, J. et al. IFNs modify the proteome of *Legionella*-containing vacuoles and restrict infection via IRG1-derived itaconic acid. PLoS Pathog 12, e1005408 (2016).

34 Schmölders, J. et al. Comparative proteomics of purified pathogen vacuoles correlates intracellular replication of *Legionella pneumophila* with the small GTPase Ras-related protein 1 (Rap1). Mol Cell Proteomics 16, 622–641 (2017).

35 Rothmeier, E. et al. Activation of Ran GTPase by a *Legionella* effector promotes microtubule polymerization, pathogen vacuole motility and infection. PLoS Pathog 9, e1003598 (2013).

36 Swart, A. L. et al. Divergent evolution of *Legionella* RCC1 repeat effectors defines the range of Ran GTPase cycle targets. mBio 11, e00405–20 (2020).

37 Steiner, B. et al. ER remodeling by the large GTPase atlastin promotes vacuolar growth of *Legionella pneumophila*. EMBO Rep 18, 1817–1836 (2017).

38 Hüsler, D., et al. *Dictyostelium* lacking the single atlastin homolog Sey1 shows aberrant ER architecture, proteolytic processes and expansion of the *Legionella*-containing vacuole. Cell Microbiol 23, e13318 (2021).

39 Hüsler, D. et al. The large GTPase Sey1/atlastin mediates lipid droplet-and FadL-dependent intracellular fatty acid metabolism of *Legionella pneumophila*. eLife 12, e85142 (2023).

40 Hedstrom, L. IMP dehydrogenase: structure, mechanism, and inhibition. Chem Rev 109, 2903–2928 (2009).

41 Naffouje, R. et al. Anti-tumor potential of IMP dehydrogenase inhibitors: a century-long story. Cancers (Basel) 11, doi:10.3390/cancers11091346 (2019).

42 Burrell, A. L. & Kollman, J. M. IMPDH dysregulation in disease: a mini review. Biochem Soc Trans 50, 71–82 (2022).

43 Natsumeda, Y. et al. Two distinct cDNAs for human IMP dehydrogenase. J Biol Chem 265, 5292–5295 (1990).

44 Senda, M. & Natsumeda, Y. Tissue-differential expression of two distinct genes for human IMP dehydrogenase (E.C.1.1.1.205). Life Sci 54, 1917-1926 (1994).

45 Collart, F. R., Chubb, C. B., Mirkin, B. L. & Huberman, E. Increased inosine-5’-phosphate dehydrogenase gene expression in solid tumor tissues and tumor cell lines. Cancer Res 52, 5826–5828 (1992).

46 Nagai, M., Natsumeda, Y. & Weber, G. Proliferation-linked regulation of type II IMP dehydrogenase gene in human normal lymphocytes and HL-60 leukemic cells. Cancer Res 52, 258–261 (1992).

47 Allison, A. C. & Eugui, E. M. Mycophenolate mofetil and its mechanisms of action. Immunopharmacology 47, 85–118 (2000).

48 Johnson, M. C. & Kollman, J. M. Cryo-EM structures demonstrate human IMPDH2 filament assembly tunes allosteric regulation. eLife 9, e53243 (2020).

49 Pimkin, M. & Markham, G. D. The CBS subdomain of inosine 5’-monophosphate dehydrogenase regulates purine nucleotide turnover. Mol Microbiol 68, 342–359 (2008).

50 Fernandez-Justel, D. et al. A nucleotide-dependent conformational switch controls the polymerization of human IMP dehydrogenases to modulate their catalytic activity. J Mol Biol 431, 956–969 (2019).

51 Plana-Bonamaiso, A. et al. Post-translational regulation of retinal IMPDH1 in vivo to adjust GTP synthesis to illumination conditions. eLife 9, doi:10.7554/eLife.56418 (2020).

52 Buey, R. M., Fernandez-Justel, D., Jimenez, A. & Revuelta, J. L. The gateway to guanine nucleotides: Allosteric regulation of IMP dehydrogenases. Protein Sci 31, e4399 (2022).

53 Burrell, A. L. et al. IMPDH1 retinal variants control filament architecture to tune allosteric regulation. Nat Struct Mol Biol 29, 47–58 (2022).

54 Ji, Y., Gu, J., Makhov, A. M., Griffith, J. D. & Mitchell, B. S. Regulation of the interaction of inosine monophosphate dehydrogenase with mycophenolic acid by GTP. J Biol Chem 281, 206–212 (2006).

55 Anthony, S. A. et al. Reconstituted IMPDH polymers accommodate both catalytically active and inactive conformations. Mol Biol Cell 28, 2600–2608 (2017).

56 Gunter, J. H. et al. Characterisation of inosine monophosphate dehydrogenase expression during retinal development: differences between variants and isoforms. Int J Biochem Cell Biol 40, 1716–1728 (2008).

57 Calise, S. J. et al. Glutamine deprivation initiates reversible assembly of mammalian rods and rings. Cell Mol Life Sci 71, 2963–2973 (2014).

58 Chang, C. C., Keppeke, G. D., Sung, L. Y. & Liu, J. L. Interfilament interaction between IMPDH and CTPS cytoophidia. FEBS J 285, 3753–3768 (2018).

59 Keppeke, G. D., Andrade, L. E. C., Barcelos, D., Fernandes, M. & Landman, G. IMPDH-based cytoophidium structures as potential theranostics in cancer. Mol Ther 28, 1557–1558 (2020).

60 Chang, C. C. et al. Molecular crowding facilitates bundling of IMPDH polymers and cytoophidium formation. Cell Mol Life Sci 79, 420 (2022).

61 Keppeke, G. D. et al. IMP/GTP balance modulates cytoophidium assembly and IMPDH activity. Cell Div 13, 5 (2018).

62 Flores-Mendez, M. et al. IMPDH2 filaments protect from neurodegeneration in AMPD2 deficiency. EMBO Rep 25, 3990–4012 (2024).

63 Peng, M. et al. The IMPDH cytoophidium couples metabolism and fetal development in mice. Cell Mol Life Sci 81, 210 (2024).

64 Chang, C. C. et al. Y12C mutation disrupts IMPDH cytoophidia and alters cancer metabolism. FEBS J 292: 3676–3695 (2025).

65 Ochtrop, P., Ernst, S., Itzen, A. & Hedberg, C. Exploring the substrate scope of the bacterial phosphocholine transferase AnkX for versatile protein functionalization. Chembiochem 20, 2336–2340 (2019).

66 Mikesh, L. M. et al. The utility of ETD mass spectrometry in proteomic analysis. Biochim Biophys Acta 1764, 1811–1822 (2006).

67 O’Neill, A. G. et al. Neurodevelopmental disorder mutations in the purine biosynthetic enzyme IMPDH2 disrupt its allosteric regulation. J Biol Chem 299, 105012 (2023).

68 Thomas, E. C. et al. Different characteristics and nucleotide binding properties of inosine monophosphate dehydrogenase (IMPDH) isoforms. PLoS One 7, e51096 (2012).

69 Allgood, S. C., et al. *Legionella* effector AnkX disrupts host cell endocytic recycling in a phosphocholination-dependent manner. Front Cell Infect Microbiol 7, 397 (2017).

70 Yu, X., et al. *Legionella* effector AnkX interacts with host nuclear protein PLEKHN1. BMC microbiology 18, 5 (2018).

71 Duan, S. Y. et al. IMPDH2 promotes colorectal cancer progression through activation of the PI3K/AKT/mTOR and PI3K/AKT/FOXO1 signaling pathways. J Exp Clin Canc Res 37: 304 (2018).

72 Xu, H. et al. IMPDH2 promotes cell proliferation and epithelial-mesenchymal transition of non-small cell lung cancer by activating the Wnt/beta-catenin signaling pathway. Oncol Lett 20, 219 (2020).

73 Oesterlin, L. K., Goody, R. S. & Itzen, A. Posttranslational modifications of Rab proteins cause effective displacement of GDP dissociation inhibitor. Proc Natl Acad Sci U S A 109, 5621–5626 (2012).

74 Gavriljuk, K. et al. Unraveling the phosphocholination mechanism of the *Legionella pneumophila* enzyme AnkX. Biochemistry 55, 4375–4385 (2016).

75 Ernst, S., et al. *Legionella* effector AnkX displaces the switch II region for Rab1b phosphocholination. Sci Adv 6, eaaz8041 (2020).

76 Molofsky, A. B. & Swanson, M. S. Differentiate to thrive: lessons from the *Legionella pneumophila* life cycle. Mol Microbiol 53, 29–40 (2004).

77 Manske, C. & Hilbi, H. Metabolism of the vacuolar pathogen *Legionella* and implications for virulence. Front Cell Infect Microbiol 4, 125 (2014).

78 Striednig, B. et al. Quorum sensing governs a transmissive *Legionella* subpopulation at the pathogen vacuole periphery. EMBO Rep 22, e52972 (2021).

79 Hayward, D. et al. ANKRD9 is a metabolically-controlled regulator of IMPDH2 abundance and macro-assembly. J Biol Chem 294, 14454–14466 (2019).

80 Rother, M. et al. Combined human genome-wide RNAi and metabolite analyses identify IMPDH as a host-directed target against *Chlamydia* infection. Cell Host Microbe 23, 661–671 (2018).

81 Yu, J. et al. Combining multi-omics analysis to identify host-targeted targets for the control of *Brucella* infection. Microb Biotechnol 16, 2345–2366 (2023).

82 Leyssen, P., Balzarini, J., De Clercq, E. & Neyts, J. The predominant mechanism by which ribavirin exerts its antiviral activity in vitro against flaviviruses and paramyxoviruses is mediated by inhibition of IMP dehydrogenase. J Virol 79, 1943–1947 (2005).

83 Khan, M., Dhanwani, R., Patro, I. K., Rao, P. V. & Parida, M. M. Cellular IMPDH enzyme activity is a potential target for the inhibition of Chikungunya virus replication and virus induced apoptosis in cultured mammalian cells. Antiviral Res 89, 1–8 (2011).

84 Li, T. W. et al. SARS-CoV-2 Nsp14 protein associates with IMPDH2 and activates NF-kappaB signaling. Front Immunol 13, 1007089 (2022).

85 Shah, C. P. & Kharkar, P. S. Discovery of novel human inosine 5’-monophosphate dehydrogenase 2 (hIMPDH2) inhibitors as potential anticancer agents. Eur J Med Chem 158, 286–301 (2018).

86 Shah, C. P. & Kharkar, P. S. Newer human inosine 5’-monophosphate dehydrogenase 2 (hIMPDH2) inhibitors as potential anticancer agents. J Enzyme Inhib Med Chem 33, 972–977 (2018).

87 Yin, Y. et al. Mycophenolic acid potently inhibits rotavirus infection with a high barrier to resistance development. Antiviral Res 133, 41–49 (2016).

88 Villarroel, M. C., Hidalgo, M. & Jimeno, A. Mycophenolate mofetil: An update. Drugs Today (Barc) 45, 521–532 (2009).

89 Zhang, H. et al. Antiviral treatment for viral pneumonia: current drugs and natural compounds. Virol J 22, 62 (2025).

90 Tiaden, A. et al. The *Legionella pneumophila* response regulator LqsR promotes host cell interactions as an element of the virulence regulatory network controlled by RpoS and LetA. Cell Microbiol 9, 2903–2920 (2007).

91 Liu, H. & Naismith, J. H. An efficient one-step site-directed deletion, insertion, single and multiple-site plasmid mutagenesis protocol. BMC Biotechnol 8, 91 (2008).

92 Cox, J. & Mann, M. MaxQuant enables high peptide identification rates, individualized p.p.b.-range mass accuracies and proteome-wide protein quantification. Nat Biotechnol 26, 1367–1372 (2008).

93 Tyanova, S. & Cox, J. Perseus: A bioinformatics platform for integrative analysis of proteomics data in cancer research. Methods Mol Biol 1711, 133–148 (2018).

94 Hughes, C. S. et al. Single-pot, solid-phase-enhanced sample preparation for proteomics experiments. Nat Protoc 14, 68–85 (2019).

95 Urwyler, S., Finsel, I., Ragaz, C. & Hilbi, H. Isolation of *Legionella*-containing vacuoles by immuno-magnetic separation. Curr Protoc Cell Biol Chapter 3, Unit 3 34 (2010).

96 Derre, I. & Isberg, R. R. *Legionella pneumophila* replication vacuole formation involves rapid recruitment of proteins of the early secretory system. Infect Immun 72, 3048–3053 (2004).

97 Hochstrasser, R. et al. The structure of the *Legionella* response regulator LqsR reveals amino acids critical for phosphorylation and dimerization. Mol Microbiol 113, 1070–1084 (2020).

98 Perez-Riverol, Y. et al. The PRIDE database and related tools and resources in 2019: improving support for quantification data. Nucleic Acids Res 47, D442–D450 (2019).

99 Segal, G. & Shuman, H. A. Intracellular multiplication and human macrophage killing by *Legionella pneumophila* are inhibited by conjugal components of IncQ plasmid RSF1010. Mol Microbiol 30, 197–208 (1998).

100 Sadosky, A. B., Wiater, L. A. & Shuman, H. A. Identification of *Legionella pneumophila* genes required for growth within and killing of human macrophages. Infect Immun 61, 5361–5373 (1993).

101 Wiater, L. A., Sadosky, A. B. & Shuman, H. A. Mutagenesis of *Legionella pneumophila* using Tn*903*dll*lacZ*: identification of a growth-phase-regulated pigmentation gene. Mol Microbiol 11, 641–653 (1994).

102 Tiaden, A. et al. The autoinducer synthase LqsA and putative sensor kinase LqsS regulate phagocyte interactions, extracellular filaments and a genomic island of *Legionella pneumophila*. Environ Microbiol 12, 1243–1259 (2010).

